# Molecular mechanisms controlling the biogenesis of the TGF-β signal Vg1

**DOI:** 10.1101/2021.04.25.441333

**Authors:** P. C. Dave P. Dingal, Adam N. Carte, Tessa G. Montague, Medel B. Lim Suan, Alexander F. Schier

**Affiliations:** Department of Molecular and Cellular Biology, Harvard University, Cambridge, MA, USA; Systems, Synthetic, and Quantitative Biology Program, Harvard, University, Cambridge, MA, USA; Biozentrum, University of Basel, Basel, Switzerland; Allen Discovery Center for Cell Lineage Tracing, University of Washington, Seattle, WA, USA; Department of Bioengineering, The University of Texas at Dallas, Richardson, TX, USA

**Keywords:** Vg1, Nodal, retention, processing, zebrafish

## Abstract

The TGF-beta signals Vg1 (Dvr1/Gdf3) and Nodal form heterodimers to induce vertebrate mesendoderm. The Vg1 proprotein is a monomer retained in the endoplasmic reticulum (ER) and is processed and secreted upon heterodimerization with Nodal, but the mechanisms underlying Vg1 biogenesis are largely elusive. Here we clarify the mechanisms underlying Vg1 retention, processing, secretion and signaling, and introduce a Synthetic Processing (SynPro) system that enables the programmed cleavage of ER-resident and extracellular proteins. First, we find that Vg1 can be processed by intra- or extracellular proteases. Second, Vg1 can be processed without Nodal but requires Nodal for secretion and signaling. Third, Vg1-Nodal signaling activity requires Vg1 processing, whereas Nodal can remain unprocessed. Fourth, Vg1 employs exposed cysteines, glycosylated asparagines, and BiP chaperone-binding motifs for monomer retention in the ER. These observations suggest two mechanisms for rapid mesendoderm induction: chaperone-binding motifs help store Vg1 as an inactive but ready-to-heterodimerize monomer in the ER, and the flexibility of Vg1 processing location allows efficient generation of active heterodimers both intra- and extracellularly. These results establish SynPro as a new in vivo processing system and define molecular mechanisms and motifs that facilitate the generation of active TGF-beta heterodimers.

**Significance:** The TGF-beta family members Nodal and Vg1 are the major inducers of mesendoderm formation during vertebrate embryogenesis. We previously established that the Vg1 proprotein is retained in the endoplasmic reticulum (ER), and that Nodal and Vg1 form heterodimers to pattern the early embryo. However, the mechanisms underlying the retention, processing, secretion, and signaling of Vg1 have been unclear. We found two mechanisms that embryos use to efficiently generate active Nodal-Vg1 heterodimers: (1) Vg1 employs its chaperone-binding motifs to ensure its retention as a ready-to-heterodimerize monomer in the ER; (2) Using a newly devised Synthetic Processing (SynPro) System, we found that Vg1 must be processed for signaling to occur, but its processing location is flexible.

## Introduction

The TGF-beta signals Nodal and Vg1 (Dvr1/Gdf3) play crucial roles in vertebrate development (1, 2), including the induction of mesendoderm and the generation of left-right asymmetry (3–17). For example, secreted Vg1-Nodal heterodimers induce a gradient of signaling that patterns the embryonic mesendoderm in zebrafish (10). Vg1-Nodal heterodimers exert their effects as ligands for a receptor complex that comprises Activin serine-threonine kinase receptors and an essential co-receptor called Oep (Tdgf1/CRIPTO) (18–20). Activated ligand-receptor complexes catalyze phosphorylation of Smad2 (pSmad2), which accumulates in the nucleus to induce the expression of mesendodermal genes (21).

We previously proposed a 4-step model for how Vg1-Nodal heterodimers pattern the mesendoderm of zebrafish embryos (10): (1) Maternally ubiquitous Vg1 proprotein is retained as a monomer in the endoplasmic reticulum (ER) of embryonic cells. (2) Expression of zebrafish Nodal genes, *cyclops* (*cyc*) and squint (*sqt*), initiates at the yolk margin at ∼3 hours post-fertilization (hpf). (3) Nodal forms heterodimers with pre-existing Vg1. (4) Vg1-Nodal heterodimers are processed and secreted to activate signaling. This model explains how Nodal and Vg1 interact during early embryogenesis, but the molecular mechanisms that regulate Vg1 retention, processing, secretion and signaling have remained unclear.

Previous studies of growth factor processing and ER retention provide potential mechanisms for how Vg1 localization, processing and activity might be regulated. TGF-beta ligands are synthesized as pre-proproteins that comprise an amino-terminal signal sequence, a long prodomain, and a shorter bioactive mature domain. A conserved cysteine in the mature domain is primarily responsible for dimer formation via an intermolecular disulfide bond (22). TGF-beta prodomain processing can occur in the Golgi apparatus and on the cell surface, where proprotein convertases are found (23–26), but it is unclear if Vg1 needs to be processed intra- or extracellularly or whether Vg1 processing depends on Nodal. It is also unclear how Vg1 is retained in the ER. Vg1 has an intact prodomain when it is retained in the ER (10), raising the possibility that Vg1 cannot be secreted because the Vg1 prodomain cannot be processed (3, 4, 27). Alternatively, a suite of protein folding chaperones could potentially interact with specific motifs on the Vg1 proprotein to block its release from the ER and subsequent processing. For example, protein disulfide isomerases (PDIs) that bind to exposed cysteines facilitate the folding of nascent proproteins and promote the retention of unassembled protein complexes in the ER (28, 29). Additionally, the lectin chaperones, calnexin and calreticulin, bind to asparagine-linked glycosyl groups on nascent secreted proteins (30). Glycosylated proteins are released from the ER after trimming off N-linked glucose moieties (30, 31). Moreover, the BiP chaperone (also known as Hspa5/Grp78) aids in protein folding by binding to predominantly hydrophobic heptapeptide sequences and can also retain proteins in the ER (32–34). Vg1 has exposed cysteines and asparagines that can be glycosylated (35), but it is unclear whether Vg1 or other TGF-beta signals use these chaperone-binding motifs to ensure retention in the ER.

In this study, we investigate the molecular mechanisms that regulate Vg1 retention, processing, secretion and signaling. To control prodomain processing, we created a novel Synthetic Processing (SynPro) system that enables programmed cleavage of ER-resident and extracellular proteins. Using SynPro, we find that Vg1 can be cleaved and activated either cell-autonomously or non-cell-autonomously. Vg1 can be processed without Nodal but requires Nodal for secretion and signaling. We further show that Vg1, but not Nodal, must be processed for signaling activity. Finally, we identify several chaperone-binding motifs in the prodomain and mature domains of Vg1 that function in ER retention. These molecular mechanisms and sequence motifs control Vg1 biogenesis and contribute to the temporal and spatial specificity of mesendoderm formation.

## Results

### Creation of a synthetic processing system (SynPro)

The activity of Vg1 depends on its dimerization with Nodal (10), but where processing must occur relative to dimerization and secretion has remained unclear. For example, Vg1 might be processed before or after secretion, and processing might occur by proteases expressed in Vg1-secreting cells or in neighboring cells. To address these questions, we first set out to control processing orthogonally by creating SynPro. SynPro employs the ability of a synthetic protease to cleave a specific peptide cleavage site (Figure 1 and SI Appendix, Fig. S1). We screened several proteases for their ability to cleave short peptide sequences and to be produced in zebrafish embryos without deleterious effects. These included proteases from the tobacco etch virus (TEVp), tobacco vein mottling virus (TVMVp), human rhinovirus 3C, and enterokinase (36). We found that most commercially available proteases are toxic when produced from mRNAs injected into zebrafish embryos (SI Appendix, Fig. S1A). Thus, we synthesized zebrafish-codon-optimized mRNAs of TEVp, TVMVp and 13 additional proteases of the *Potyviridae* family (37). All 15 proteases were non-toxic when expressed in zebrafish embryos (SI Appendix, Fig. S1B).

**Figure 1.**
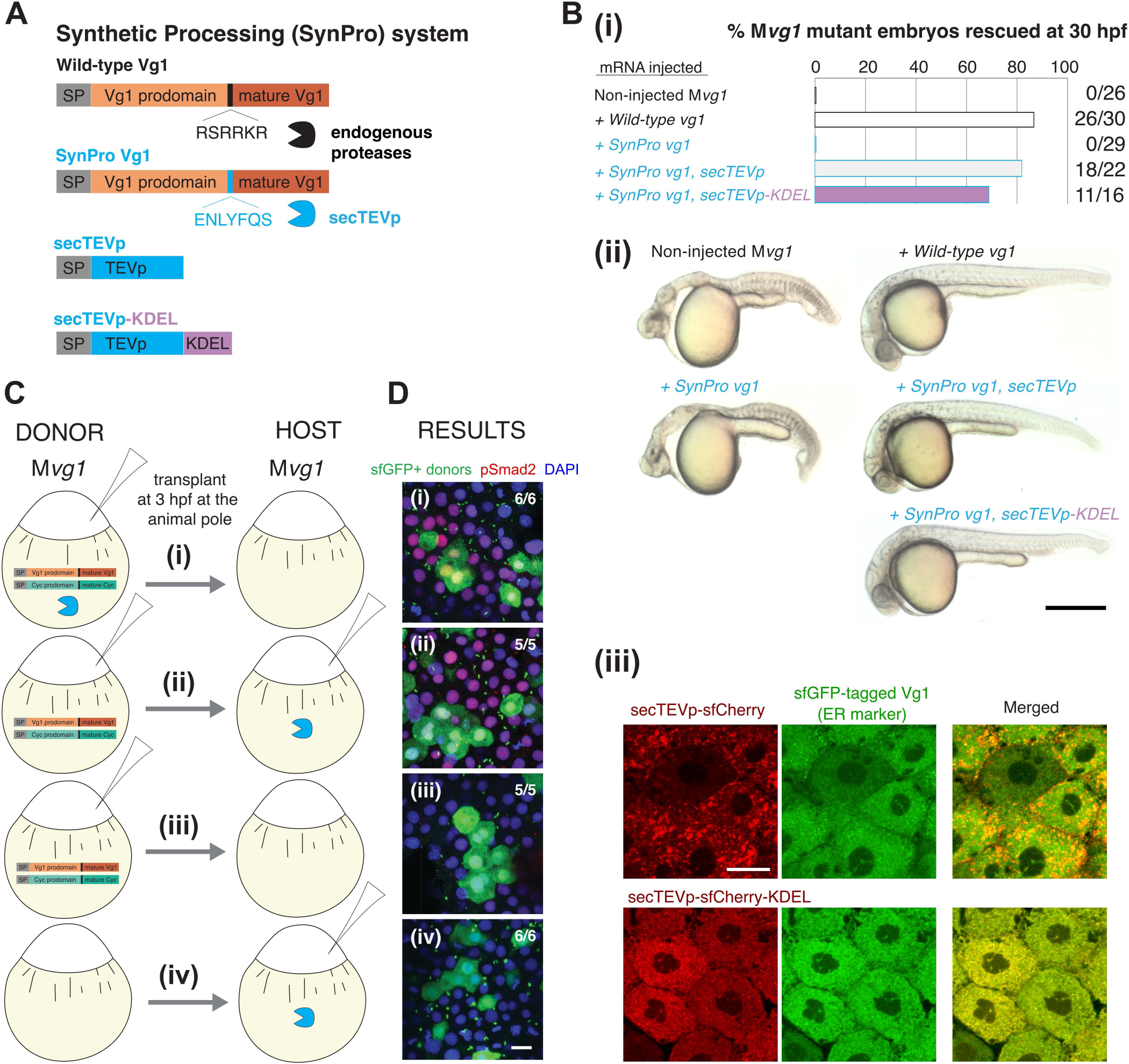
The Synthetic Processing (SynPro) system shows that prodomain cleavage can be non-cell autonomous. (**A**) The SynPro system comprises an orthogonal secreted protease derived from tobacco etch virus (secTEVp) and a cognate sequence that replaces the endogenous cleavage site of Vg1 (SynPro Vg1, RSRRKR → ENLYFQS). (**B**) (**i**) Rescue percentage after 30 hpf of M*vg1* embryos injected with 50 pg of *vg1*, *SynPro vg1*, *SynPro vg1* and *secTEVp*, or *SynPro vg1* and *secTEVp-KDEL* mRNAs. (**ii**) Representative images of 30 hpf M*vg1* embryos for the indicated injection condition. Minor brain and tail defects are noted in embryos transiently rescued with mRNAs of the SynPro system. Scale bar, 50 mm. (**iii**) Fluorescence images of M*vg1* embryos at 50-60% epiboly that were injected with mRNAs for sfGFP-tagged wild-type Vg1 and secTEVp-sfCherry (top panels) or secTEVp-sfCherry-KDEL (bottom panels). Scale bar, 20 μm. (**C**) Schematic of transplantation assay. M*vg1* embryos were injected with 50 pg mRNA each of: (**i**) DONOR: *cyc*, *SynPro vg1*, and *secTEVp*; HOST: none; (**ii**) DONOR: *cyc* and *SynPro vg1*; HOST: *secTEVp*; (**iii**) DONOR: *cyc* and *SynPro vg1*; HOST: none; (**iv**) DONOR: none; HOST: *secTEVp*. All M*vg1* donor embryos were marked by also injecting 50 pg *sfGFP* mRNA. At high stage, before the onset Nodal signaling, sfGFP-marked DONOR cells were transplanted to the animal pole of HOST M*vg1* embryos. (**D**) At 50-60% epiboly, chimeric embryos were fixed and immunostained for sfGFP and pSmad2. DAPI, nuclei. Scale bar, 20 μm.

To test whether the synthetic proteases are functional, we designed a fluorescent reporter of proteolytic cleavage (SI Appendix, Fig. S1C; see Materials and Methods for details). Briefly, a protease-cleavable substrate is initially localized in the cytoplasm. Protease-catalyzed cleavage results in the release into the nucleus and reconstitution of a split fluorescent protein, mNeonGreen2 (38). Co-expression of the cleavable substrate and the nuclear reporter in zebrafish embryos did not lead to reconstitution of nuclear mNeonGreen2 fluorescence (SI Appendix, Fig. S1D,i). By contrast, nuclear mNeonGreen2 fluorescence was observed when TEVp was co-expressed with its cognate substrate and the nuclear reporter (SI Appendix, Fig. S1D,ii). We observed protease-induced fluorescence reconstitution for 6 out of 15 proteases tested (SI Appendix, Fig. S1E), indicating that the synthetic proteases can cleave their cognate sequences in zebrafish embryos.

### Synthetically processed Vg1 rescues *vg1* mutants

While the TEV protease has been shown to be active in the cytoplasm, none have been shown to be active in the secretory system *in vivo*. To generate a secreted protease for the SynPro system, we added an amino-terminal signal sequence to TEVp to produce a secreted variant, secTEVp (see Materials and Methods for details). We also introduced five amino acid substitutions in secTEVp that are known to promote solubility (39) and prevent oxidation in the secretory compartments (40). To determine whether secTEVp is functional and can replace endogenous convertases that cleave the Vg1 prodomain, we generated SynPro Vg1. In this Vg1 derivative, the native cleavage sequence, ‘RSRRKR’, was replaced with the cognate cleavage sequence of secTEVp, ‘ENLYFQS’ (Figure 1A). Remarkably, the co-expression of *SynPro vg1* and *secTEVp* rescued mesendoderm formation in maternal *vg1* mutant (M*vg1*) embryos, whereas the expression of *SynPro vg1* mRNA alone failed to rescue M*vg1* mutants (Figure 1B). Notably, co-production of SynPro Vg1 and an ER-localized secTEVp-KDEL (Figure 1B,iii) also rescued M*vg1* embryos, revealing that intracellular processing of Vg1 can result in normal Vg1 activity. These results show that endogenous enzymes that process Vg1 can be replaced by an orthogonal protease and that SynPro provides a novel tool to control processing and protein activity in the secretory system.

### Vg1 processing is not sufficient for secretion

The co-expression of *SynPro vg1* and *secTEVp* rescued M*vg1* mutants but did not induce abnormal overexpression phenotypes. This result suggests that Vg1 signaling activity was restricted to domains of co-expression with endogenous Nodal, raising two hypotheses: (1) Vg1 cleavage by SynPro depends on Nodal or (2) Vg1 cleavage by SynPro is independent of Nodal, but its secretion requires Nodal. To test these models, we co-expressed *SynPro vg1* and ER-resident *secTEVp* and assessed the location and processing of SynPro Vg1 with or without the zebrafish Nodal *cyc* (SI Appendix, Fig. S2). We found that SynPro Vg1 was processed but not secreted in the absence of the Nodal signal Cyclops. These results support hypothesis (2): synthetic processing of Vg1 does not require Nodal, but Nodal is necessary for Vg1 secretion.

### Vg1 processing can be non-cell autonomous

The results above show that intracellular processing in the secretory compartments is sufficient to produce functional Vg1. We next determined whether processing of the Vg1 proprotein in the extracellular milieu might also be sufficient to generate active Vg1. By leveraging the versatility of the SynPro system, we generated scenarios where SynPro Vg1 and secTEVp were expressed in the same or in different cells. We performed a series of transplant experiments in M*vg1* embryos that were injected with mRNAs expressing components of the SynPro system (Figure 1C-D). First, donor M*vg1* cells that co-express *SynPro vg1*, *secTEVp* and *cyc* mRNAs were transplanted into host M*vg1* embryos (Figure 1C). In this scenario, ligands and protease are co-expressed in the same cells. As expected, we observed Nodal signaling activity based on immunostaining of nuclear pSmad2 in both donor and neighboring host cells (Figure 1D,i). This result indicates that donor cells secreted active secTEVp-processed SynPro Vg1 and Cyc. Second, donor cells co-expressing *SynPro vg1* and *cyc* were transplanted into host M*vg1* embryos expressing *secTEVp*. In this scenario, ligands and protease are not co-expressed in the same cells. Strikingly, we observed nuclear pSmad2 accumulation in both donor and surrounding host cells (Figure 1D,ii). In control experiments, nuclear pSmad2 was never observed when secTEVp was absent, nor when donor embryos did not express SynPro Vg1 and Cyc (Figure 1D,iii; Figure 1D,iv). Thus, host-secreted secTEVp was able and required to cleave and activate the donor-secreted SynPro Vg1 and Cyc. This application of the SynPro system indicates that Vg1 can be processed and activated non-cell autonomously.

### Processing is not required for secretion of Vg1 and Nodal

The observation that extracellular secTEVp was sufficient to generate active signaling around cells producing SynPro Vg1 and Nodal suggested that an unprocessed Vg1 proprotein can be secreted in the presence of Nodal. To further test this idea, we generated non-cleavable mutants of Vg1 and Nodal. We inactivated the cleavage site of Vg1 and of the zebrafish Nodal Squint (Sqt) and inserted a superfolder green fluorescent protein (sfGFP) to generate the non-cleavable variants, *vg1-NC-sfGFP* and *sqt-NC-sfGFP*. We could not generate a non-cleavable variant of Cyc due to additional cryptic cleavage sites (SI Appendix, Fig. S3). We found that both sfGFP-tagged Vg1 (*vg1-sfGFP*) or non-cleavable Vg1 (*vg1-NC-sfGFP*) were secreted when co-produced with Sqt or non-cleavable Sqt (*sqt-NC*) in M*vg1* embryos (Figure 2A). Notably, endogenous levels of Cyc-Vg1 and Sqt-Vg1 heterodimers are so low that secretion of Vg1-sfGFP cannot be detected without ectopic addition of Sqt (10). Secretion was incomplete under conditions in which at least one signal was non-cleavable, as evidenced by intracellular and cell membrane localization. These results indicate that Vg1 and Nodal can be secreted in the absence of processing.

**Figure 2.**
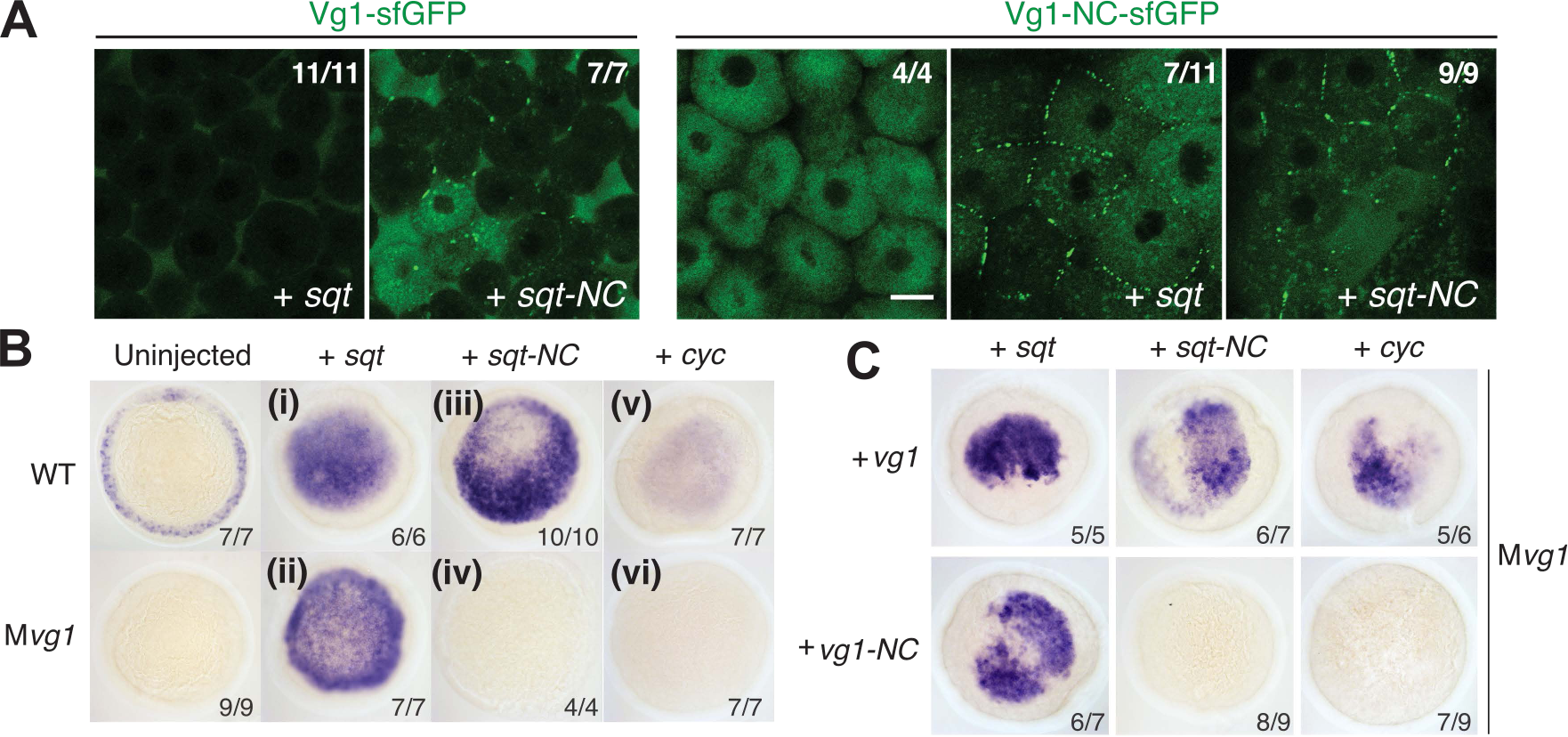
Prodomain cleavage affects Vg1-Nodal signaling but not secretion. (**A**) Live fluorescence imaging of M*vg1* co-injected with 50 pg of *vg1-sfGFP* or non-cleavable *vg1-sfGFP* (*vg1-NC-sfGFP*, RSRRKR → SQNTSN) mRNA and 50 pg of *sqt* or *sqt-NC* (RRHRR → SQNTS) mRNA. Scale bar, 17 μm. (**B**) Nodal target gene (*lefty1*) expression at 50% epiboly in WT and M*vg1* embryos injected with 50 pg of *sqt*, *sqt-NC* or *cyc* mRNA. (**C**) *lefty1* expression in M*vg1* embryos co-injected with 50 pg *sqt*, *sqt-NC* or *cyc* and *vg1* or *vg1-NC* mRNA.

### Processing is not required for Nodal activity in the presence of processed Vg1

To test whether non-cleavable Vg1 (Vg1-NC) and Nodal are physiologically active, we injected mRNAs of the non-cleavable constructs into wild-type and M*vg1* embryos and assessed the induction of Nodal target genes. Remarkably, non-cleavable Sqt was able to induce target gene expression in embryos expressing wild-type Vg1 (Figure 2B,iii, iv). By contrast, non-cleavable Sqt was inactive in the absence of Vg1 (M*vg1* mutants) or when co-expressed with Vg1-NC (Figure 2C). Vg1-NC was also inactive when co-expressed with Cyclops (Figure 2B,v-vi). These results indicate that the Vg1 prodomain – but not the Nodal prodomain – must be cleaved for active heterodimer signaling.

### Cysteine thiol and N-glycosyl groups retain the Vg1 prodomain in the ER

Previous studies have shown that replacement of the Vg1 prodomain with other TGF-beta prodomains resulted in Vg1 processing and mesoderm-inducing activity (3, 4, 27, 41, 42). Similarly, we found that a chimeric Vg1 protein that fused the Nodal prodomain to the Vg1 mature domain resulted in a secreted and active Vg1 (SI Appendix, Fig. S4A). Conversely, replacing the prodomain of zebrafish Nodals with the prodomain of Vg1 inhibited the secretion and activity of these chimeric TGF-betas (SI Appendix, Fig. S4B). These results suggest that the Vg1 prodomain has features that block secretion and promote ER retention.

To determine how the Vg1 prodomain might mediate ER retention, we searched for putative sequence motifs that might mediate retention. We did not find any KDEL sequences in Vg1; these motifs are found at the C-termini of ER-resident proteins and are recognized by KDEL receptors that trigger Golgi-to-ER retrograde transport (43). However, we found putative sequence motifs in Vg1 that bind to ER-resident chaperones, including exposed cysteines that bind to PDIs (28, 29) and glycosylated asparagines in NX[S/T] motifs that bind to calnexin and calreticulin (30). In particular, the prodomain of Vg1 (proVg1) has one exposed cysteine (C100) and two potential N-glycosylation sites (N108, N179). Notably, the mature domain of Vg1 (matVg1) also has a free cysteine (C319) and a potential N-glycosylation site (N296), raising the possibility that the mature domain also plays a role in ER retention (Figure 3A). To determine whether these residues are accessible at the surface, we used AlphaFold2 (44) (there is no biophysically determined Vg1 structure). The top model for Vg1 in AlphaFold2 shows that the cysteine and asparagine residues are indeed exposed at the surface (Figure 3B), suggesting that ER-resident chaperones can potentially access them.

**Figure 3.**
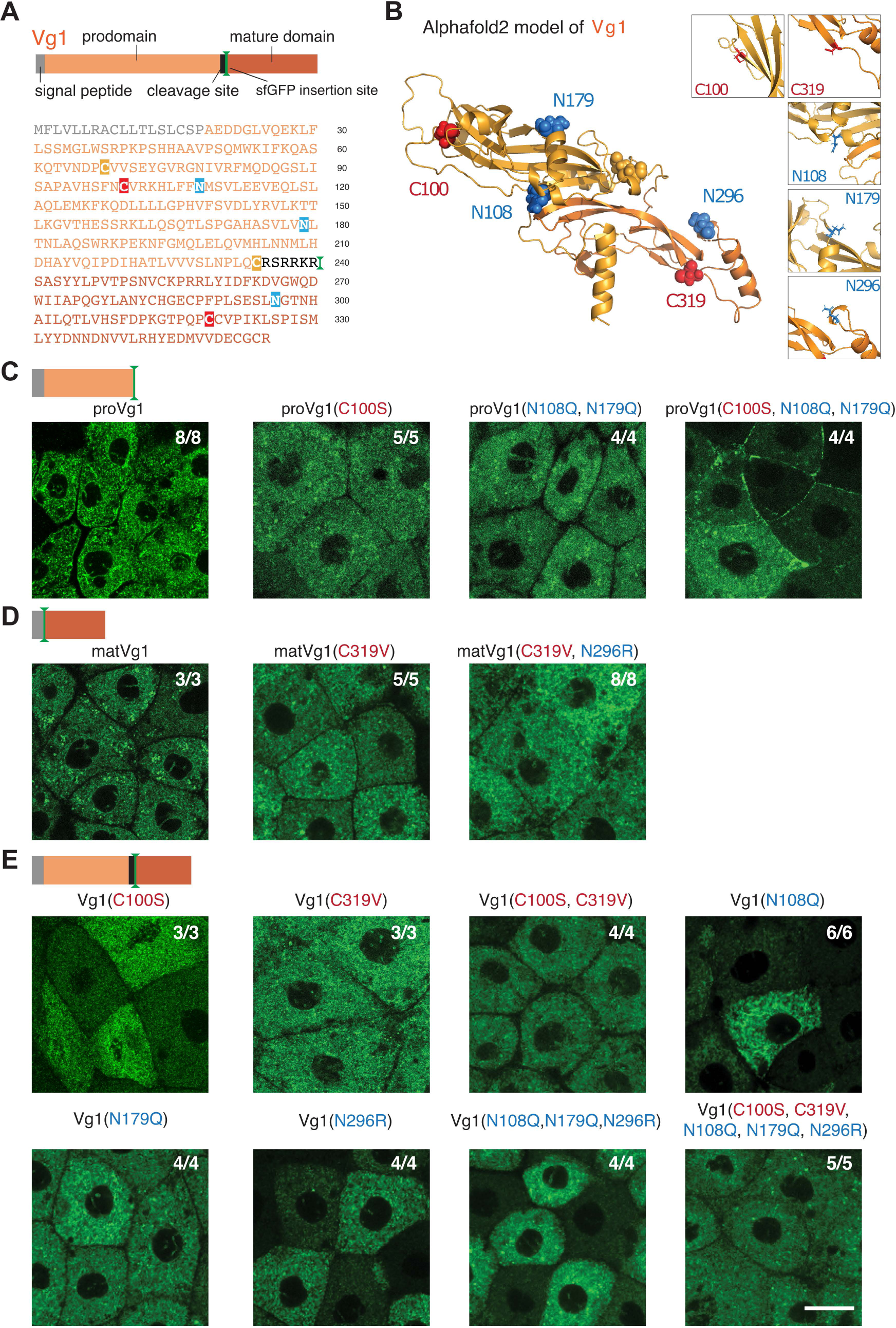
Cysteine and N-linked glycosylation sites retain the Vg1 prodomain in the ER. (**A**) Schematic and primary amino acid sequence of zebrafish Vg1 pre-proprotein. Cysteines (red, yellow) and asparagines (blue) are highlighted. (**B**) Alphafold2 model for Vg1 is shown in cartoon representation, whereas the cysteine (red, yellow) and asparagine (blue) residues are shown in spheres (insets show zoomed-in views of the residues mutated in this study). We make a minor note here that two cysteine residues, C68 and C234 (yellow), are predicted to form a disulfide bond and thus were not further studied. (**C-E**) Fluorescence images of fixed M*vg1* embryos injected with 50 pg mRNA of *sfGFP*-tagged *vg1* prodomain (proVg1) (**C**), *vg1* mature domain (matVg1) (**D**), and full-length *vg1* (**E**), with or without the indicated cysteines and asparagines mutated. *sfGFP* was inserted into *vg1* downstream of the predicted basic cleavage site in all constructs. Scale bar, 20 μm.

To elucidate whether these residues promote Vg1 retention in the ER, we systematically mutated the cysteine and N-glycosylation residues in sfGFP-tagged Vg1, proVg1, and matVg1 constructs and visualized their localization. We injected M*vg1* embryos with *sfGFP*-tagged *vg1*, *proVg1*, or *matVg1* mRNAs and performed fluorescence microscopy to determine the subcellular localization of the resulting proteins. Similar to full-length Vg1, we observed ER localization of sfGFP-tagged proVg1 and matVg1 (Figure 3C-D). Mutants for the prodomain cysteine (C100S) or glycosylation sites (N108Q, N179Q) were retained in the ER, but the triple mutant proVg1(C100S, N108Q, N179Q) was secreted to the extracellular space (Figure 3C). By contrast, the matVg1 cysteine and glycosylation mutants (C319S, N296R) as well as the quintuple mutant Vg1(C100S, N108Q, N179Q, C319S, N296R) were still retained in the ER (Figure 3D, E). We verified the secretion or localization of these constructs in the ER of M*vg1* embryos by co-expressing them with a fluorescent ER marker (sfCherry-KDEL) or with a nucleocytoplasmic marker (sfCherry-Smad2) (SI Appendix, Fig. S4C). We also found that the loss of N-glycosylation sites was more deleterious to signaling activity than the loss of cysteines (SI Appendix, Fig. S5). Taken together, our mutagenesis results show that N-linked glycosyl groups and an exposed cysteine thiol retain the Vg1 prodomain in the ER.

### BiP-binding motifs retain the Vg1 mature domain in the ER

The observation that the free cysteine and N-glycosylation mutants of matVg1 and full-length Vg1 are still retained in the ER suggests that matVg1 possesses additional ER-retention motifs. We hypothesized that a third chaperone, BiP (Hspa5/Grp78), might promote ER retention. To elucidate which sequence features of matVg1 may bind to BiP, we utilized the Gething-Sambrook scoring system (34). Since mature Nodals (matSqt and matCyc) are secreted (SI Appendix, Fig. S6), we compared the sequences of matVg1 and matCyc and scored for all possible BiP-binding heptapeptides (Figure 4A-B, Table 1; see Materials and Methods for details). To test whether the high-scoring heptapeptides of matVg1 promote ER retention, we systematically mutated three high-scoring matVg1 sequences to their corresponding low-scoring matCyc sequences (labeled as *m1*, *m2*, and *m3* in Figure 4A, C). While all single mutants and double mutants were found in the ER, the triple mutant matVg1(*m1*, *m2*, *m3*) exhibited extracellular localization (Figure 4D). Conversely, a mutant matCyc possessing the high-scoring features of matVg1 was retained in the ER (Table 1 and SI Appendix, Fig. S7). These results indicate that BiP-binding sequences retain the Vg1 mature domain in the ER.

**Figure 4.**
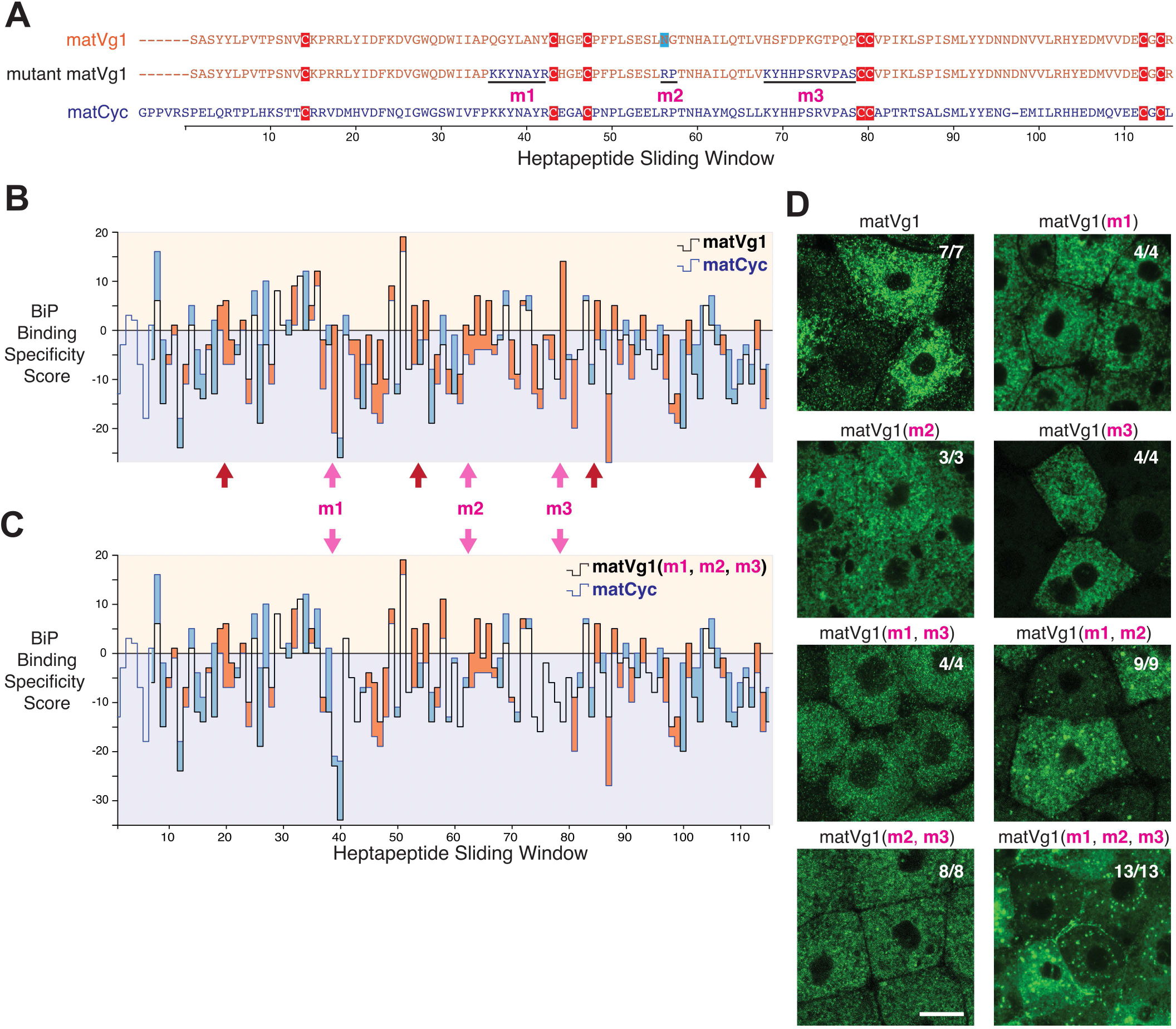
Binding motifs for BiP promote ER retention of the Vg1 mature domain. (**A**) Amino acid sequence alignment of the mature domains of Vg1 (matVg1) and Cyc (matCyc) and mutant matVg1. Cysteines (red) and asparagines (blue) are highlighted. (**B-C**) Difference charts of the BiP binding specificity scores (34) between two protein sequences along a sliding window of seven amino acids. Orange fills indicate the matVg1 or matVg1(*m1*, *m2*, *m3*) score (black line) > matCyc score (blue line). Conversely, blue fills indicate matCyc score > matVg1 or matVg1(*m1*, *m2*, *m3*) score. Arrows denote regions where matVg1 score > 0 and matCyc score < 0. Red arrows specifically denote regions that contain cysteines involved in cystine-knot formation. (**B**) Differences in BiP binding specificity scores between matVg1 (black line) and matCyc (blue line). (**C**) Differences in BiP binding specificity scores between mutant matVg1(*m1*, *m2*, *m3*) (black line) and matCyc (blue line). Note the loss of orange fills in *m1*, *m2*, and *m3* regions (pink arrows) when compared to (**B**). (**D**) Fluorescence images of fixed M*vg1* embryos injected with 50 pg mRNA of sfGFP-tagged *vg1* mature domain variants. *sfGFP* was inserted upstream of the *vg1* mature domain in all constructs.

**Table 1.**
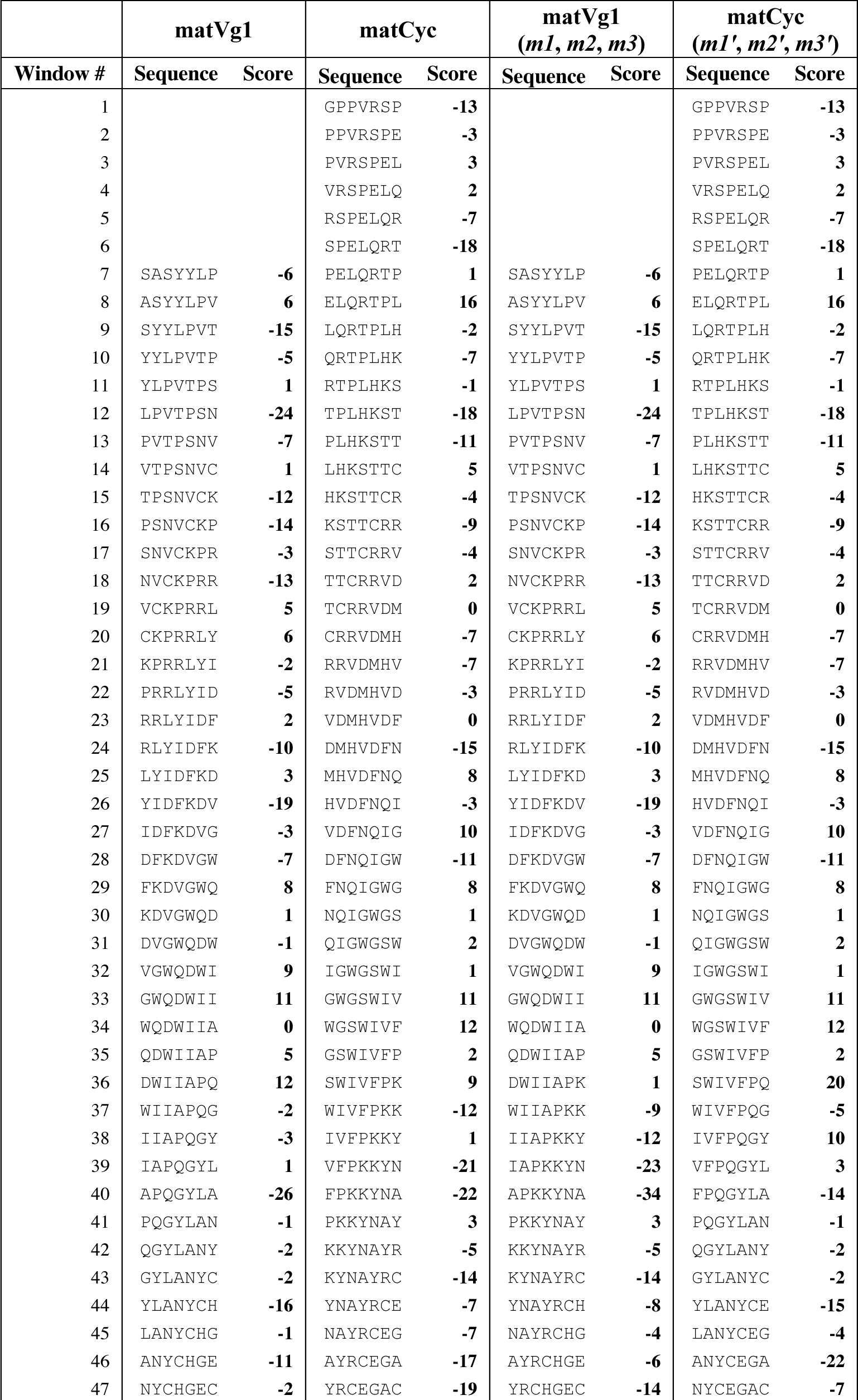
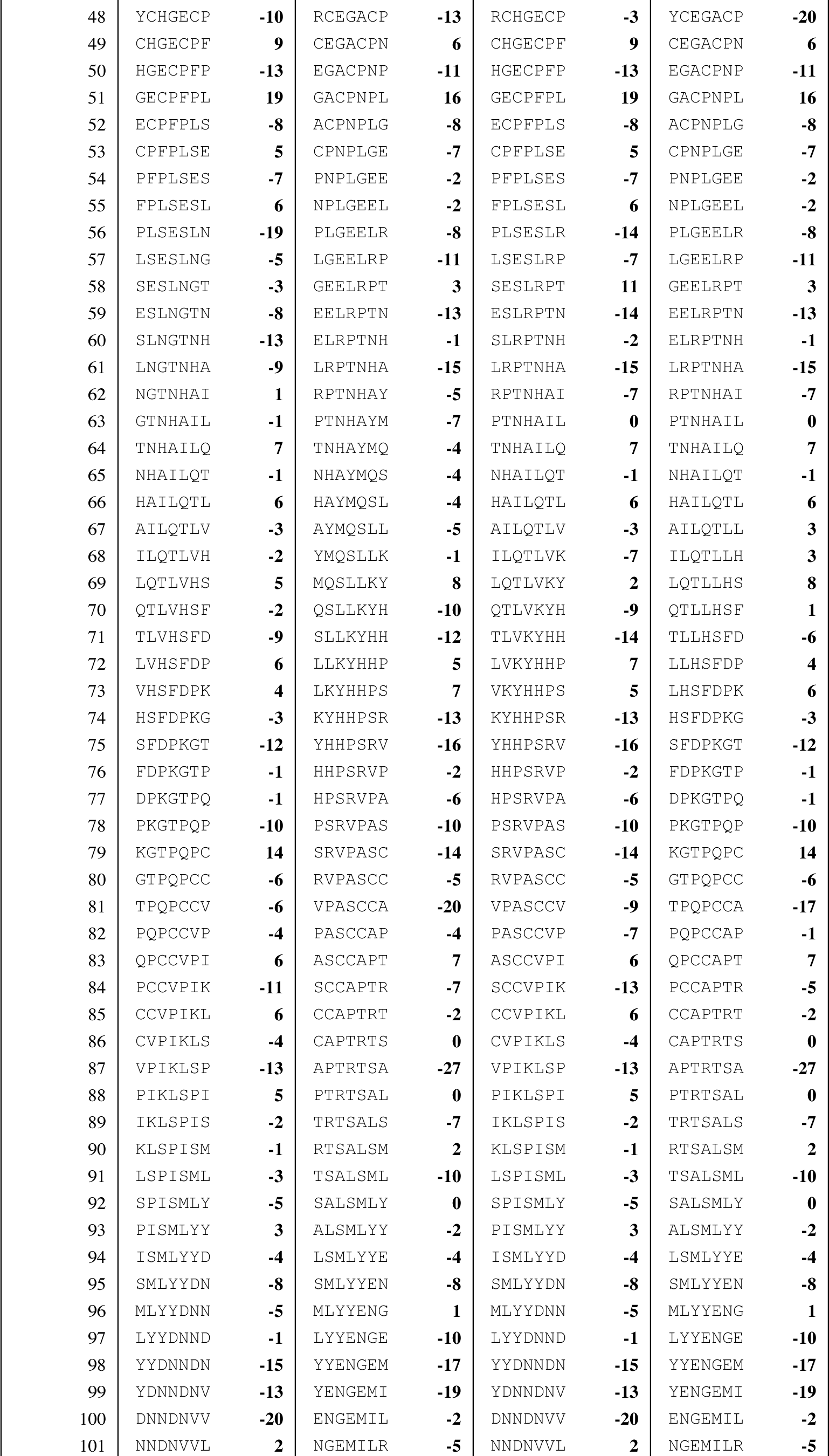
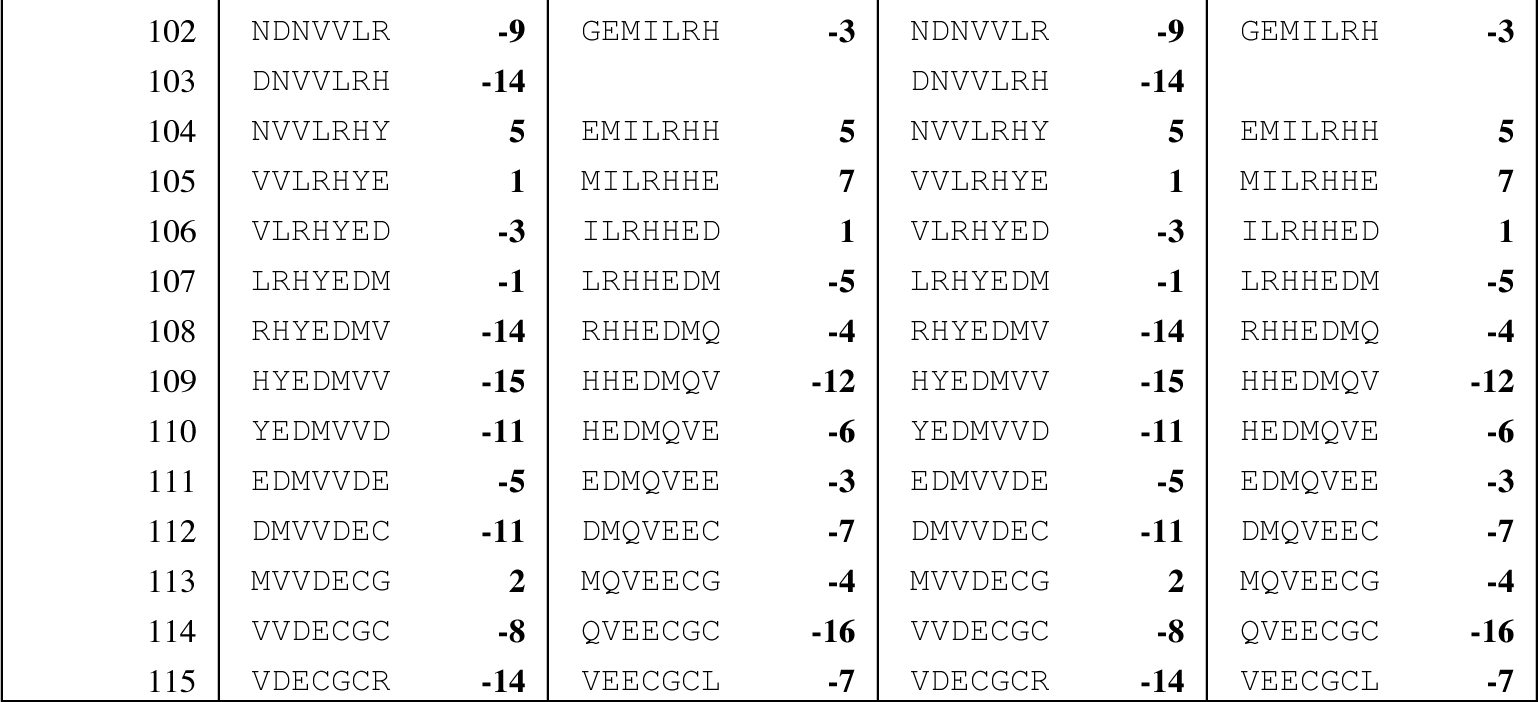
BiP Binding Specificity Scores of matVg1, matCyc, matVg1(*m1*, *m2*, *m3*), and matCyc(m1’, m2’, m3’) sequences. These scores are visualized in Figure 4B-C and in SI Appendix, Fig. S7.

## Discussion

This study reveals features of Vg1 that regulate its retention, processing, secretion, and signaling during early zebrafish embryogenesis (Figure 5): (1) Vg1 ER retention is mediated by exposed cysteines, glycosylated asparagines, and BiP chaperone-binding motifs. (2) Vg1 can be processed without Nodal but requires Nodal for secretion and signaling. (3) Vg1 can be processed cell-autonomously or non-cell autonomously. (4) Vg1-Nodal signaling activity requires processing of Vg1 but not of Nodal. These conclusions unify and extend several previous observations about Vg1-Nodal signaling:

**Figure 5.**
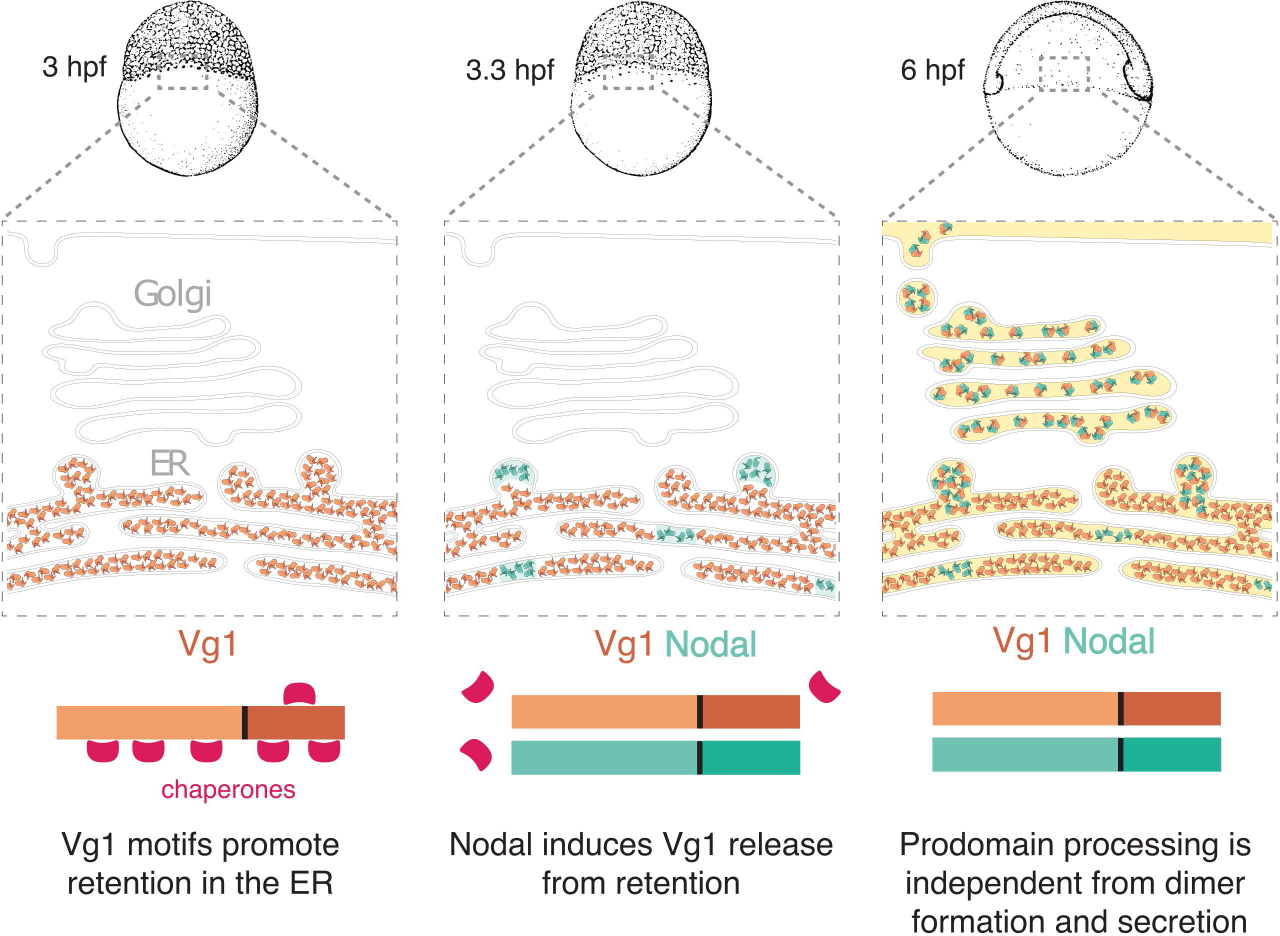
Model of Vg1-Nodal heterodimer formation and processing. Maternal Vg1 is retained in the ER via chaperone-binding motifs; at the maternal-zygotic transition, Nodal is produced and induces release of Vg1 from the ER via heterodimer formation; processing of the Vg1 prodomain is required for the activity of Vg1-Nodal heterodimers but can be independent from dimer formation and secretion.

First, our study defines Vg1 sequence motifs that have the hallmarks of binding sites for ER-resident chaperones. Our mutational analyses indicate that these motifs help retain monomeric Vg1 in the ER until Nodal is expressed and heterodimerizes with Vg1. We find that the Vg1 prodomain is specifically retained in the ER via its N-glycosylation sites and exposed cysteine. Thus, Vg1 is a member of a class of secreted proteins that are transiently retained in the ER until their cysteines form intermolecular disulfide bonds (45–48). In addition, Vg1 employs BiP-binding regions as chaperone motifs for ER retention. Mutating the BiP-binding regions (*m1*, *m2*, *m3*) of the Vg1 mature domain leads to secretion. Interestingly, the *m3* region is juxtaposed to cysteine C319, which mediates heterodimerization with Nodal. It is conceivable that BiP binding blocks the formation of mature Vg1 homodimers. We note that Vg1 mutant constructs that remain in the ER may have signaled to ER chaperones that they are misfolded, preventing us from determining whether additional retention motifs exist or whether mutations in the construct led to misfolding.

We found no evidence for KDEL-mediated ER retention of Vg1. This mode of ER retention differs from chaperone-mediated retention in that it mediates retrograde transport from the Golgi apparatus to the ER lumen (43). By contrast, chaperone-mediated retention prevents the escape of proteins from the ER. We speculate that the Vg1 system has evolved to trap Vg1 in the ER until Nodal is imported and displaces the chaperones as a heteromeric partner.

Second, our study reveals shared features of the Vg1-Nodal system with the heavy- and light-chain immunoglobulin (Ig) assembly system (46, 49–60). Both Nodal and free light chains and are readily secreted (57, 58), whereas both Vg1 and heavy chains are retained in the ER until they are covalently bonded to Nodal and light chains, respectively. Mutating the exposed cysteine in the heavy chain leads to premature secretion of monomers (59, 60). Our results suggest a similar role for the exposed cysteine in Vg1, but in addition, Vg1 also employs N-glycosylation sites and BiP-binding regions as chaperone motifs for ER retention. During antibody assembly, the BiP chaperone can stably bind to unstructured heavy chains (55, 59) and cooperates with PDI and calnexin to properly assemble light and heavy chains (52, 61). A similar mechanism might control Vg1-Nodal heterodimer assembly. In this scenario, the Vg1 monomer remains in an unstructured, immature state through interaction with various chaperones in the ER and might fold only upon interaction with Nodal. How the chaperone network hands over the Vg1 monomer to Nodal remains to be elucidated.

Third, our study clarifies the roles of Vg1 processing in secretion. Early studies suggested that Vg1 cannot be secreted because its prodomain cannot be processed (3, 4, 27). However, our application of the SynPro system shows that prodomain processing in the ER is not sufficient to induce the secretion of Vg1 (SI Appendix, Fig. S2). In fact, unprocessed Vg1 can be secreted upon dimerizing with Nodal (Figure 2). These results are consistent with the observations that mouse Nodal and its proprotein convertases are not co-expressed in the same cell (24, 25, 62, 63). In this case, prodomain processing of Gdf1/3 (the orthologs of Vg1) and Nodal dimers occurs after they are released in the extracellular space or bound to the target cell surface and in early endosomes. Using the SynPro system in zebrafish embryos, we show that Vg1 can also be cleaved after its Nodal-induced release in the extracellular space (Figure 1). These findings establish that prodomain processing of Vg1 can be location-independent and separable from Nodal-induced secretion.

Fourth, our results elucidate the roles of Nodal and Vg1 processing in signaling. We find that Vg1-Nodal heterodimers with non-cleavable prodomains are secreted but cannot signal, whereas unprocessed Nodal in combination with processed Vg1 remains partially active. This result demonstrates that Nodal processing is not required for secretion or partial activity, consistent with previous studies that found that non-processed mouse Nodal still promotes mesoderm formation but cannot position or maintain the primitive streak (64). Our finding that the Vg1 prodomain must be processed for signaling suggests the possibility that the activity of non-processed mouse Nodal also relies on processed Vg1 (Gdf1/3).

Fifth, our study extends the previous suggestion that the Vg1-Nodal heterodimeric system allows for the rapid activation of the Nodal signaling pathway (10). Instead of a time delay caused by the accumulation of sufficient levels of Nodal needed to form homodimers, the preformed pool of Vg1 allows for rapid heterodimer formation as soon as Nodal is synthesized. Our study suggests that two features of Vg1 facilitate this strategy of rapid signaling: chaperone-mediated monomer retention in the ER stores Vg1 in an inactive but ready-to-heterodimerize form, and location-independent processing allows efficient generation of active heterodimers both intra- and extracellularly.

Last, the SynPro system introduced here has broad applications in the targeted processing and regulation of secreted proteins. Our results show that endogenous Vg1 can be replaced by a Vg1 derivative containing an engineered cleavage sequence in combination with the corresponding SynPro protease. The M*vg1* rescue experiments with the exogenous SynPro system exhibited minor morphological defects, which might be more fully recovered when the SynPro system is stably integrated into the genome. The SynPro system might be used for the orthogonal regulation of TGF-beta and other secreted signals. Endogenous proprotein convertases cleave the same polybasic motif in multiple secreted proteins. By contrast, each SynPro protease recognizes a unique cleavage sequence. Different SynPro proteases could be combined to separately control the activity of multiple secreted signals. For example, different TGF-beta signals could be processed independently to reveal distinct spatial and temporal requirements. Beyond TGF-beta proteins, the SynPro system could be used to independently process several bioactive peptides from hormone or neuropeptide precursors. By releasing individual peptides one at a time from a polyprotein, the effects of each peptide could be analyzed. Thus, the SynPro system has the potential to accelerate the functional assignment of bioactive peptides generated by secreted polyproteins.

## MATERIALS AND METHODS

### Genotyping of *vg1* mutants

Genomic DNA was isolated via the HotSHOT method from either excised adult caudal fin tissue or individual fixed embryos (65). Genotyping was carried out via PCR using standard conditions followed by 2% gel electrophoresis. Mutant *vg1* fish have the *vg1^a165^* allele, which contains a 29 bp deletion in the first exon of *vg1* and was detected as described (10). Allele designation was omitted for brevity in the rest of the text.

### Zebrafish husbandry and fish lines

Fish were maintained per standard laboratory conditions (66). Embryos were raised at 28.5°C in embryo medium (250 mg/L Instant Ocean salt, 1 mg/L methylene blue in reverse osmosis water adjusted to pH 7 with sodium bicarbonate) and staged according to a standard staging series (67). Wild-type fish and embryos represent the TLAB strain. The *vg1* mutant fish line was maintained as previously described (10). M*vg1* embryos were generated by crossing zygotic homozygous *vg1* female fish to TLAB wild-type male fish.

### Cloning of expression constructs and the Synthetic Processing (SynPro) system

Standard molecular cloning techniques, such as PCR, Gibson Assembly (68), and site-directed mutagenesis, were performed to assemble new constructs used in this study.

The coding sequences (CDS) of *vg1* (10), *sqt*, *cyc* (69), and their variants tagged with superfolder GFP (sfGFP) (70), were previously assembled into the pCS2(+) vector that contains a β-globin 5’ UTR and an SV40 late polyA signal at the 3’ UTR. For sfGFP-tagged Vg1 prodomain or mature domain, site-directed mutagenesis (Q5 Kit, New England Biolabs) of pCS2(+)-vg1-sfGFP was performed to truncate the full-length construct into respective domains (as depicted in Figure 3). All point mutants, indels, and epitope tags were subsequently generated using site-directed mutagenesis. For non-cleavable vg1-NC, vg1-NC-sfGFP and proVg1-sfGFP, the cleavage site ‘RSRRKR’ was replaced with ‘SQNTSN’. For non-cleavable sqt-NC and sqt-NC-sfGFP, the cleavage site ‘RRHRR’ was replaced with ‘SQNTS’. For chimeras of Squint/Cyclops prodomain and sfGFP-tagged Vg1 mature domain (sfGFP-matVg1), the Squint and Cyclops prodomains (including their respective cleavage sites) were PCR-amplified from pCS2(+)-squint and pCS2(+)-cyclops, and sfGFP-matVg1 was PCR-amplified from pCS2(+)-vg1-sfGFP.

Commercially available sequences for tobacco etch virus protease (TEVp, Addgene Plasmid #8835), tobacco vein mottling virus protease (TVMVp; Addgene Plasmid #8832), and human rhinovirus 3C protease (HRV 3Cp; Addgene Plasmid #78571) were gifts from David Waugh (71–73). Commercially available sequence for bovine enterokinase (Addgene Plasmid #49048) was a gift from Hans Brandstetter (74). These sequences were PCR-amplified and subcloned into the pCS2(+) vector for downstream applications *in vivo*.

Zebrafish codon-optimized sequences of 15 Potyviral proteases listed in Supp Fig 3 were synthesized by Integrated DNA Technologies and subcloned into the pCS2(+) vector. For the protease cleavage reporter assay described in Supp Fig 3, Component 1 is a cleavable substrate for Potyviral proteases and was constructed via Gibson assembly of zebrafish beta-Arrestin (*arrb2a* gene PCR-amplified from high-stage cDNA library) and mNeonGreen2(1–10) (a gift from Bo Huang) (38) into the pCS2(+) vector. The cognate cleavage site inserted between arrb2a and mNeonGreen2(1–10) was encoded in the primer overhangs. Component 2 is the codon-optimized Potyviral protease described above. Component 3 was constructed via Gibson assembly of four tandem copies of mNeonGreen2(11) (a gift from Bo Huang) (38) and mCherry-tagged Histone 2B (a gift from Jeffrey Farrell) into the pCS2(+) vector.

For the SynPro system, the codon-optimized TEV protease was further modified using site-directed mutagenesis to generate secTEVp. The secTEVp construct contains a secretion signal sequence derived from zebrafish toddler (translated signal peptide: MRFFHPLYLLLLLLTVLVLISA) and 5 point mutations (N23Q, C130S, T173G, L56V, S135G). The secTEVp-sfCherry-KDEL construct additionally contains a C-terminal fusion of sfCherry3C (38) sequence and an ER-targeting motif ‘(GSGS)EEKDEL’. SynPro Vg1 was derived from pCS2(+)-vg1-sfGFP using site-directed mutagenesis to replace the ‘RSRRKR’ cleavage site to ‘(GSGS)ENLYFQS(GS)’. The ER marker sfCherry-KDEL was generated using site-directed mutagenesis of the sfCherry3C sequence to insert the secretion signal sequence from toddler at the N-terminus and the C-terminal sequence ‘EEKDEL’. To fluorescently mark the cytoplasm and nucleus, sfCherry-Smad2 was cloned via Gibson assembly, where sfCherry3C is N-terminal to zebrafish Smad2 via a ‘GSGSGS’ linker. To verify whether the zebrafish proteome contain sequences identical to the cleavage motif of secTEVp, a BLASTP alignment search showed that a secreted protein RgmA contains the best sequence match of ‘EDLYFQS’, which is predicted by Alphafold2 to be not surface-accessible (https://alphafold.ebi.ac.uk/entry/Q1LVM8) and should also not be recognized by secTEVp.

### Determination of BiP-binding scores

To focus our mutagenesis efforts, we ignored corresponding matVg1 and matCyc heptapeptides that both scored positive in the Gething-Sambrook scoring system because BiP is predicted to bind to both. We also ignored corresponding regions where both heptapeptides scored zero or negative because BiP is predicted to not bind at all. We focused on 8 regions wherein matVg1 heptapeptides scored positive and the corresponding matCyc sequences scored negative (Figure 2B, arrows). We ignored 4 out of the 7 regions because they included six cysteines that participate in cystine-knot formation (Figure 2B, red arrows). BiP-binding scores were visualized using the D3.js chart library (https://d3js.org/) and the code used is available on GitHub (https://github.com/davedingal/BiP_binding_score).

### mRNA synthesis and microinjection into embryos

All pCS2(+) plasmids were linearized with NotI and subsequently purified with the E.Z.NA. Cycle Pure Kit (Omega). Capped mRNAs were synthesized with the Sp6 mMessage Machine kit (Invitrogen) using the purified linearized plasmids as templates. Capped mRNAs were then purified with the E.Z.N.A. Total RNA Kit I (Omega). Capped mRNA concentrations were measured using the NanoDrop 2000 spectrophotometer (Thermo Fisher Scientific). All kits were used according to respective manufacturer’s protocols. If not mentioned otherwise, all mRNAs were injected at 50 pg into embryos at the one-cell stage using standard methods (66).

### Transplantation

For transplantation experiments, donor and host M*vg1* embryos at one-cell stage were injected with 50 pg of each mRNA relevant to the experiment and then grown to 1000-cell stage (3 hpf). At 1000-cell stage, cells were transplanted from donor embryos to host embryos, and were grown to shield stage (6 hpf) before fixation for immunostaining.

### Live imaging and immunofluorescence imaging

Embryos were injected with 50 pg of each mRNA, grown to sphere stage, and then embedded in 1% low melting temperature agarose (Aquapor) on glass-bottomed dishes (MatTek). Imaging was performed on a Zeiss LSM 880 inverted confocal microscope.

### *in situ* hybridization

Embryos were grown to 50% epiboly then fixed in 4% formaldehyde overnight at 4°C. Whole mount *in situ* hybridizations were performed according to standard protocols (75). A DIG-labeled antisense RNA probe targeting *lefty1* was synthesized using a DIG Probe Synthesis Kit (Roche). NBT/BCIP/Alkaline phosphatase-stained embryos were dehydrated in methanol before clearing and imaging in 2:1 benzyl benzoate:benzyl alcohol (BBBA) using a Zeiss Axio Imager.Z1 microscope.

### Immunoblotting

Embryos were injected at the 1-cell stage with 50 pg of mRNA then allowed to develop to 50% epiboly. 8 embryos per sample were manually deyolked with forceps and frozen in liquid nitrogen. Samples were boiled for 5 min at 95°C with 2x SDS loading buffer and DTT (150 mM final concentration), then loaded onto Any kD protein gels (Bio-Rad). Samples were transferred to PVDF membranes (GE Healthcare), blocked in 5% non-fat milk (Bio-Rad) in TBST and incubated in primary antibodies (1:5000 rabbit anti-GFP, ThermoFisher A11122, RRID:<u>AB_221569</u>) at 4°C overnight. Proteins were detected using HRP-coupled secondary antibody (1:15,000 goat anti-rabbit, Jackson ImmunoResearch Labs 111-035-144, RRID:<u>AB_2307391</u>). Chemiluminescence was detected using Amersham ECL reagent (GE Healthcare).

### Immunostaining

A previous protocol (76) was modified to improve signal-to-noise ratio. Embryos were fixed in 4% paraformaldehyde overnight at 4°C in PBSTw (1× phosphate-buffered saline + 0.1% v/v Tween 20), washed three times in PBSTw for 10 min each, dehydrated in a MeOH/PBSTw mixture series (25%, 50%, 75%, and 100% methanol) at 5 min per wash at room temperature (RT), and stored in 100% MeOH at – 20°C for at least 2 h. Embryos were rehydrated in a MeOH/PBSTr (1× PBS + 1% Triton X-100) mixture series (75%, 50%, 25% MeOH) for 5 min each, washed three times in PBSTr for 10 min each at RT, and manually de-yolked. Embryos were then incubated in antibody binding buffer (PBSTr + 1% v/v dimethyl sulfoxide) for 2 h at RT and subsequently stored overnight at 4°C in antibody binding buffer containing relevant primary antibodies. After primary antibody incubation, embryos were washed six times with PBSTr for 10 min each, before a 30-min incubation in antibody binding buffer at RT. Embryos were then incubated for 2 h at RT in antibody binding buffer containing appropriate fluorescent secondary antibodies. Embryos were then washed six times with PBSTr. To label DNA in cell nuclei, embryos were incubated with 1 μg/mL DAPI in PBSTw for 30 min at RT. Lastly, embryos were washed three times in PBSTw for 10 min each at RT, and then mounted for microscopy.

Primary antibodies were used against GFP (1:1000 chicken IgY, Aves Lab, RRID:<u>AB_2307313</u>) and phosphorylated Smad2 (1:1000 rabbit IgG, Cell Signaling, RRID:<u>AB_2798798</u>). Fluorescent secondary antibodies used were goat anti-chicken Alexa 488 conjugate (1:2000, Thermo Fisher, RRID:<u>AB_2534096</u>) and goat anti-rabbit Alexa 647 conjugate (1:2000, Thermo Fisher, RRID:<u>AB_2633282</u>).

### Image processing

All images were processed in Fiji/ImageJ (77). Brightness, contrast, and color balance were uniformly applied to images.

## Acknowledgements

The authors thank Richard Losick, Nathan Lord, Maxwell Shafer, Madalena Madeira Reimão Pinto, Jan Christian and two reviewers for helpful comments on the manuscript, Yiqun Wang for the figure schematics, and Stephen D. Hansen for programming and data visualization. We thank the members of the Schier lab for advice, expertise, insights, and discussions. We also thank the Harvard Center for Biological Imaging and Jacopo Ferruzzi for microscopy infrastructure and support. This project was supported by NIH DP1-HD094764, the Allen Discovery Center for Cell Lineage Tracing, and a UT Dallas Startup Fund to Dr. Dingal.

## Ethics

Animal experimentation: All vertebrate animal works was performed at the facilities of Harvard University, Faculty of Arts & Science (HU/FAS). The HU/FAS animal care and use program maintains full AAALAC accreditation, is assured with OLAW (A3593-01) and is currently registered with the USDA. This study was approved by the HU/FAS Standing Committee on the Use of Animals in Research & Teaching under Protocol No. 25-08.

## Supporting Information

**Fig. S1.**
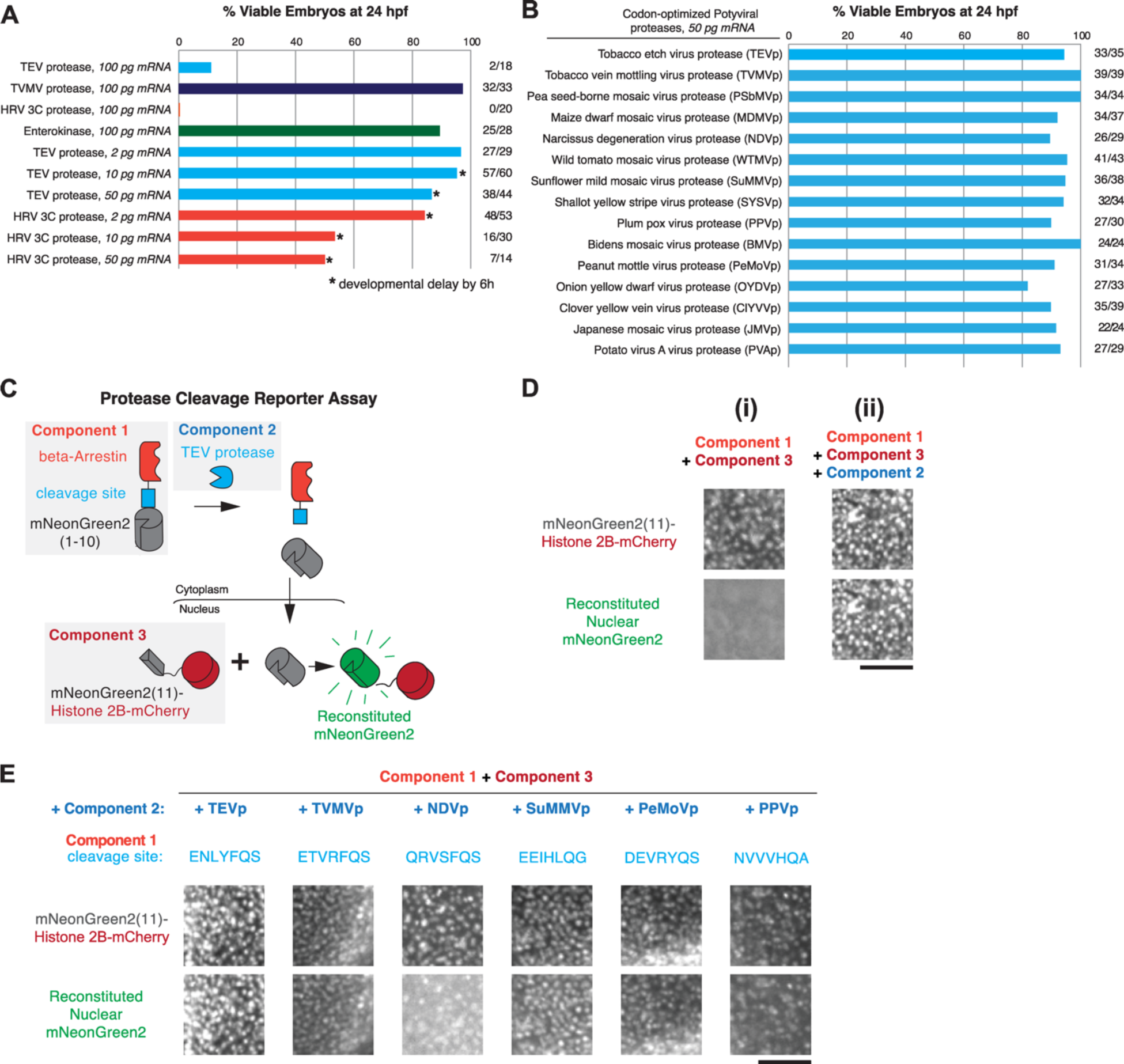
Developing the SynPro system. (**A**) Viability at 24 hpf of wild-type embryos injected with mRNA of commercially available proteases: tobacco etch virus (TEV) protease (Addgene plasmid #8835), tobacco vein mottling virus (TVMV) protease (Addgene plasmid #8832), enterokinase (Addgene plasmid #49048), and human rhinovirus 3C (HRV 3C) protease (Addgene plasmid #78571). Developmental delays and loss of viability were observed in embryos injected with TEV protease and HRV 3C protease. (**B**) Viability at 24 hpf of wild-type embryos injected with 50 pg of zebrafish codon-optimized mRNA sequences of 15 potyviral proteases. In all conditions, more than 80% of embryos were viable at 24 hpf. (**C**) Schematic of fluorescence reporter assay for protease cleavage, which is composed of three components: Component 1 contains the cleavage site flanked by beta-Arrestin and mNeonGreen2(1–10); Component 2 is the codon-optimized Potyviral protease (e.g. TEV protease recognizes the sequence ENLYFQS); and Component 3 is nuclear histone 2b tagged with mCherry and fused with mNeonGreen2(11). See Methods for construct details. (**D**) Testing the cleavage efficiency of TEV protease. (**i**) Co-expression of Components 1 and 3 does not exhibit mNeonGreen2 fluorescence, whereas (**ii**) co-expression of all 3 components leads to proteolytic cleavage, nuclear translocation, and binding of mNeonGreen2(1–10) to mNeonGreen2(11) – leading to reconstitution of mNeonGreen2 fluorescence in the nucleus. (**E**) Using the reporter assay in zebrafish embryos, six out of fifteen Potyviral proteases exhibited proteolytic activity, based on reconstitution of nuclear mNeonGreen2 fluorescence (bottom row images). Scale bars, 100 μm.

**Fig. S2.**
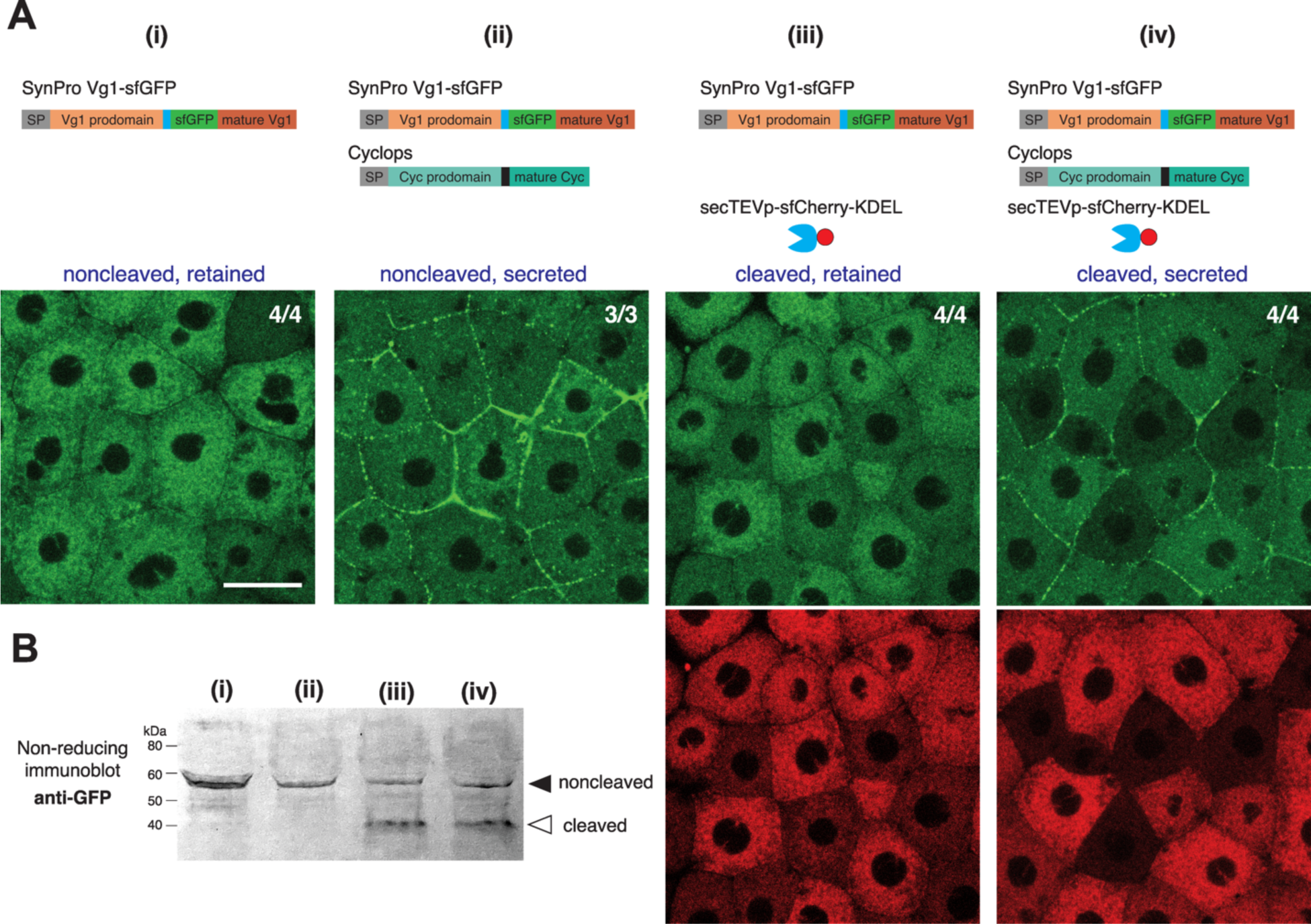
Vg1 prodomain processing does not promote secretion. (**A**) Fluorescence images of fixed M*vg1* embryos injected with 25 pg mRNA of sfGFP-tagged *SynPro vg1*, *secTEVp-sfCherry-KDEL*, and *cyc* at the indicated combinations. *sfGFP* was inserted upstream of the *vg1* mature domain. Scale bar, 20 μm. (**B**) Anti-GFP non-reducing immunoblot of M*vg1* embryos injected with the indicated constructs as in (**A**). Black arrowhead indicates the position of uncleaved SynPro Vg1, open arrowhead indicates secTEVp-cleaved Vg1. 15 embryos at 3 hpf were loaded per well.

**Fig. S3.**
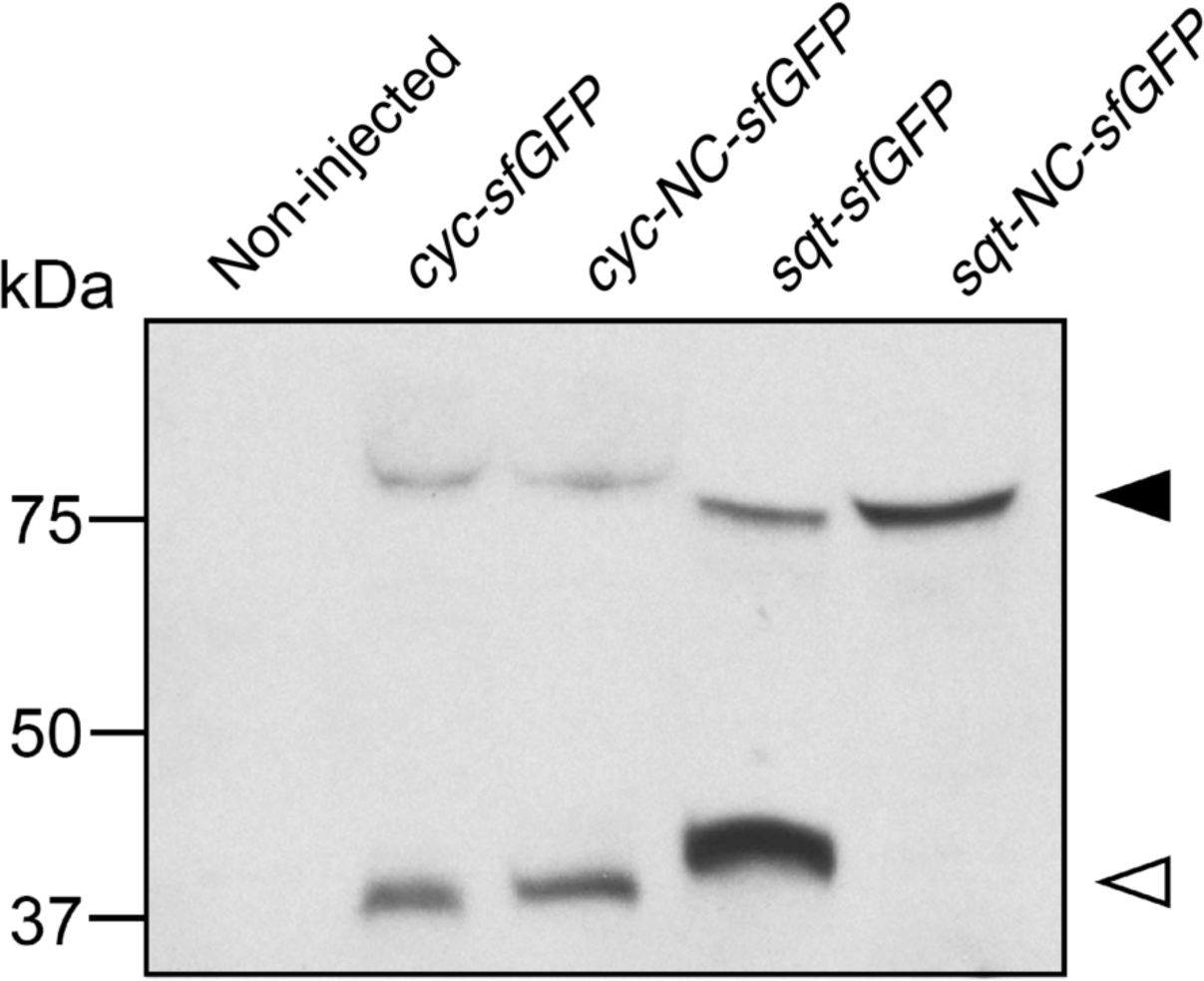
Non-cleavable Squint. Anti-GFP reducing immunoblot of M*vg1* embryos injected with 50 pg mRNA of *cyc-sfGFP*, *cyc-NC-sfGFP* (RRGRR → SQNTS), *sqt-sfGFP* or *sqt-NC-sfGFP* mRNA. Black arrowhead indicates the position of full-length protein, open arrowhead indicates processed mature domain.

**Fig. S4.**
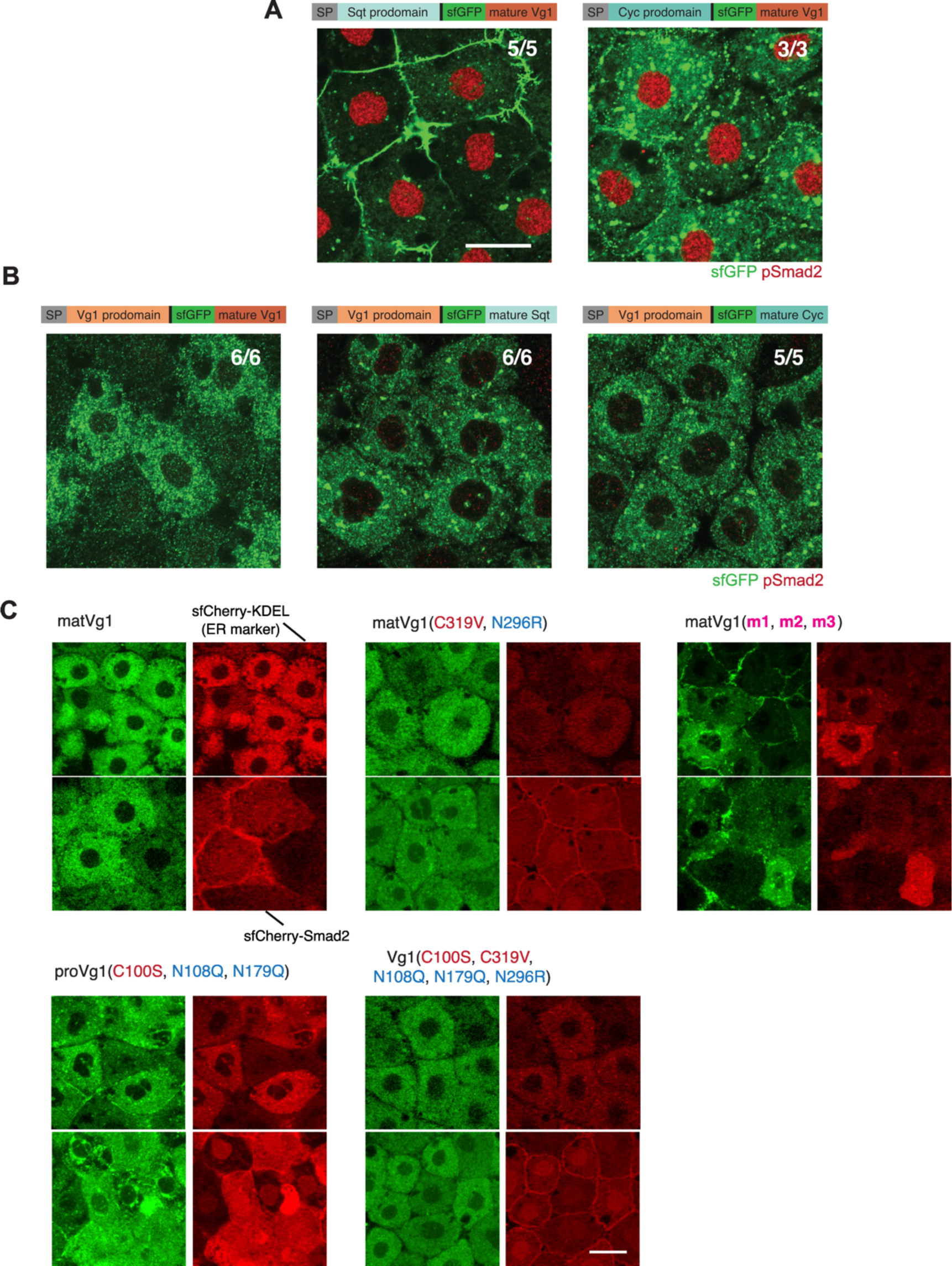
Localization of Vg1 and Nodal variants. **(A)** Immunofluorescence images of fixed M*vg1* embryos injected with 50 pg mRNA of *sqt* prodomain (left) or *cyc* prodomain (right) fused to *sfGFP*-tagged mature *vg1* domain; or injected with 50 pg mRNA of (**B**) *vg1-sfGFP* (left), *vg1* prodomain fused to *sfGFP*-tagged mature *sqt* domain (middle) or fused to *sfGFP*-tagged mature *cyc* domain (right). Embryos were immunostained with antibodies against sfGFP and pSmad2. (**C**) Fluorescence images of fixed M*vg1* embryos to verify secretion or localization in the ER of several Vg1 constructs. sfCherry-KDEL was used to mark the ER compartment; whereas sfCherry-Smad2 was used to mark the inner leaflet of the plasma membrane, cytoplasm, and nucleus (see Materials and Methods for construct details). Embryos were injected with 50 pg mRNA per construct. Scale bars, 20 μm.

**Fig. S5.**
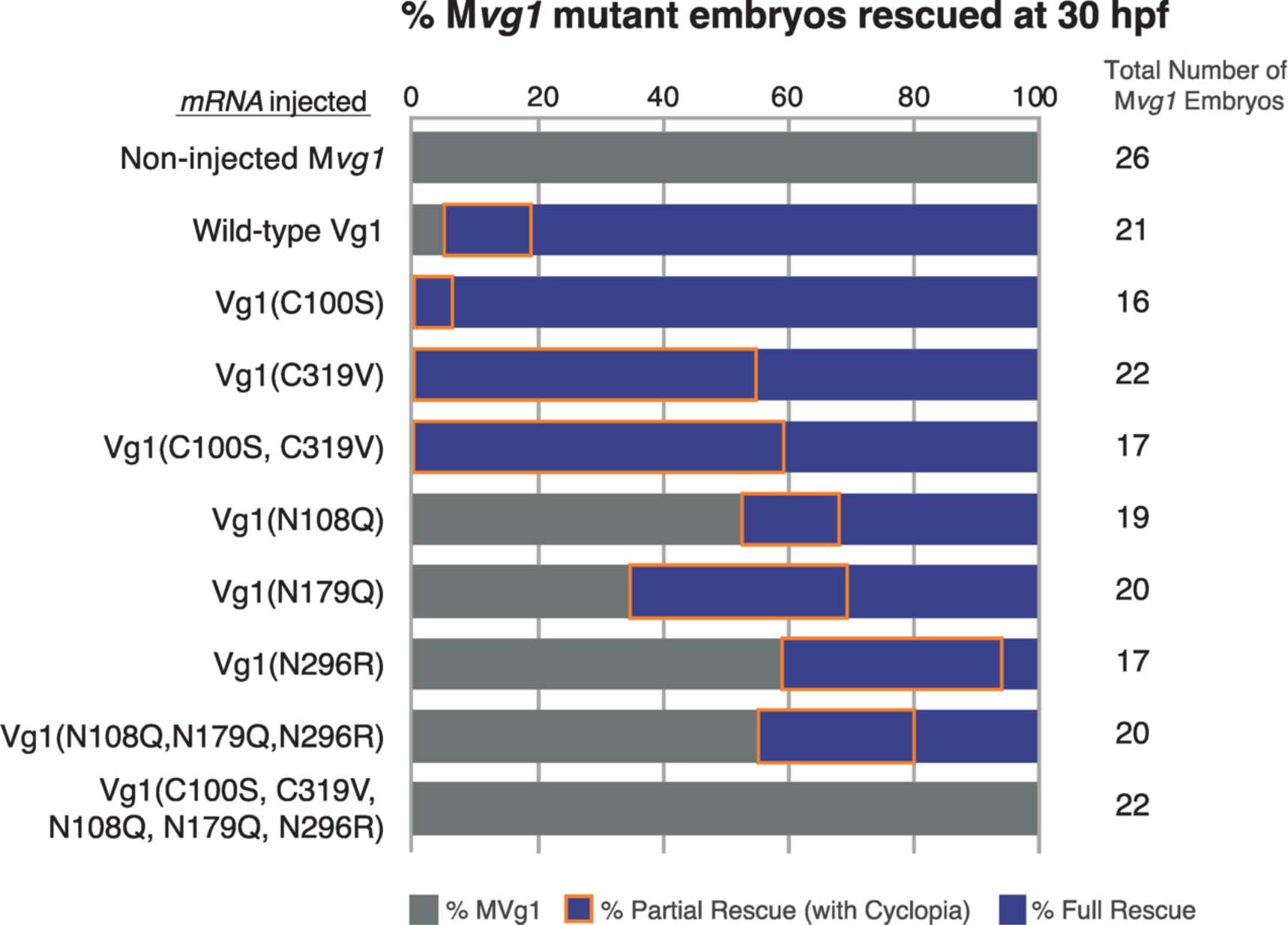
Vg1 mRNA rescue of M*vg1* embryos. Rescue percentage after 30 hpf of M*vg1* embryos injected with 50 pg of *vg1* mRNAs with the indicated mutations. Full or partial (with cyclopia) rescue percentages are respectively indicated as bars that are blue or blue with orange edges.

**Fig. S6.**
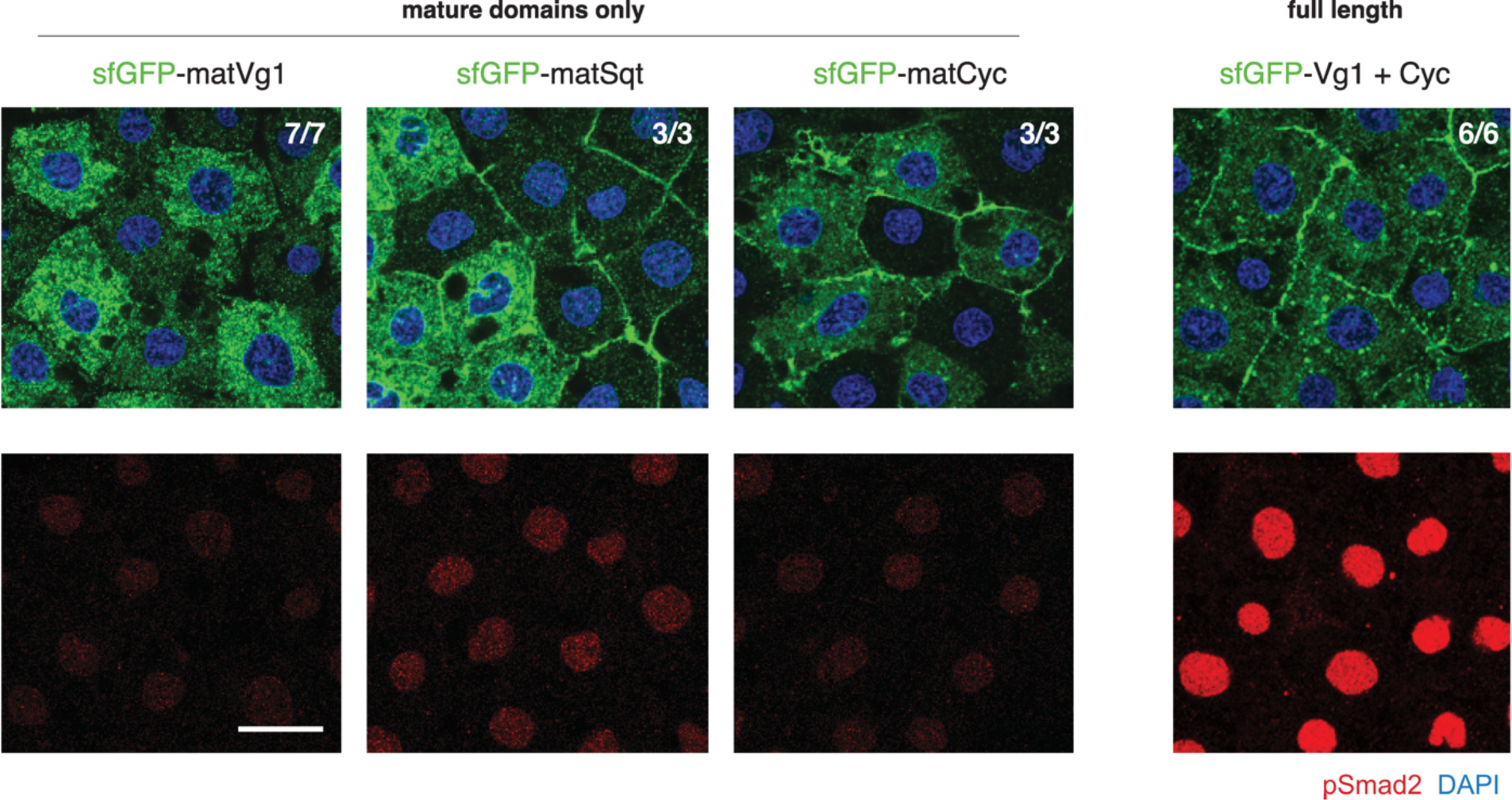
Localization and activity of Vg1 and Nodal mature domains. **(A)** (Left to right) Immunofluorescence images of fixed M*vg1* embryos injected with 50 pg mRNA of *sfGFP*-*matVg1*, *sfGFP-matSqt*, *sfGFP-matCyc*, or *sfGFP*-*matVg1* and *cyc*. The observed fluorescence in the ER for matCyc and matSqt conditions may be a result of delayed or inefficient release of the overexpressed proteins. Embryos were immunostained with antibodies against sfGFP and pSmad2. DAPI, nuclei. Scale bar, 20 μm.

**Fig. S7.**
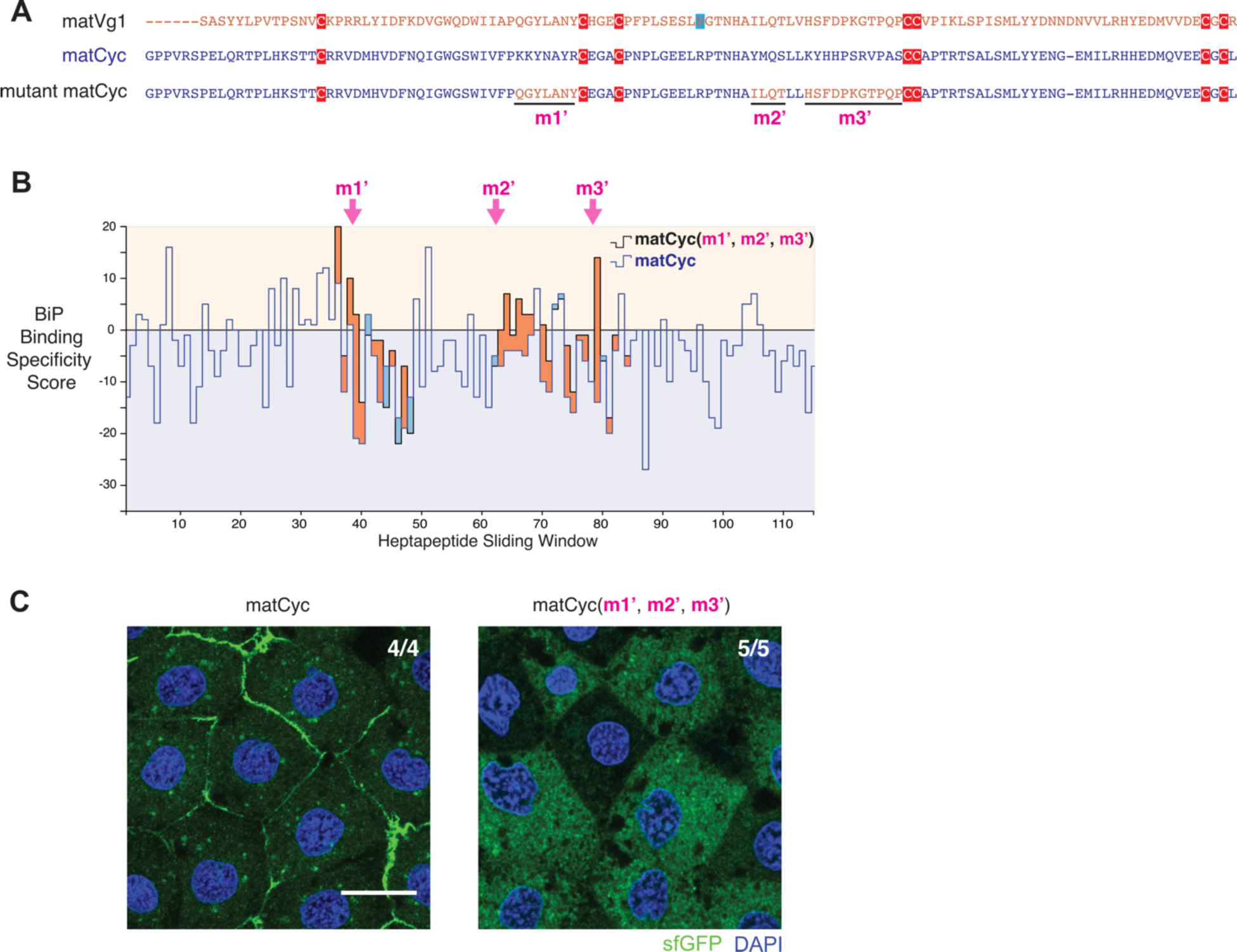
ER retention of matCyc using BiP-binding features of matVg1. (**A**) Amino acid sequence alignment of the mature domains of Vg1 (matVg1) and Cyc (matCyc) and mutant matCyc. Cysteines (red) and asparagines (blue) are highlighted. (**B**) Difference charts of the BiP binding specificity scores (34) between wild-type and mutant matCyc sequences along a sliding window of seven amino acids. Orange fills indicate the mutant matCyc (black line) score > matCyc (blue line) score. Conversely, blue fills indicate matCyc score > mutant matCyc score. Note the gain of orange fills in *m1’*, *m2’*, and *m3’* regions (arrows). (**C**) Fluorescence images of fixed M*vg1* embryos injected with 50 pg mRNA of *sfGFP*-tagged *matCyc* and *sfGFP*-tagged mutant *matCyc(m1’, m2’, m3’)*. *sfGFP* was inserted upstream of the mature *cyc* domain in all constructs. DAPI, nuclei. Scale bar, 20 μm.

## References

1. Schier AF. Nodal morphogens. Cold Spring Harb Perspect Biol. 2009;1(5):a003459. Epub 2010/01/13. doi: 10.1101/cshperspect.a003459. PubMed PMID: 20066122; PubMed Central PMCID: PMC2773646.

2. Rogers KW, Muller P. Nodal and BMP dispersal during early zebrafish development. Dev Biol. 2019;447(1):14–23. Epub 2018/04/14. doi: 10.1016/j.ydbio.2018.04.002. PubMed PMID: 29653088.

3. Dale L, Matthews G, Colman A. Secretion and mesoderm-inducing activity of the TGF-beta-related domain of Xenopus Vg1. EMBO J. 1993;12(12):4471–80. Epub 1993/12/01. PubMed PMID: 8223457; PubMed Central PMCID: PMC413871.

4. Thomsen GH, Melton DA. Processed Vg1 protein is an axial mesoderm inducer in Xenopus. Cell. 1993;74(3):433–41. Epub 1993/08/13. doi: 10.1016/0092-8674(93)80045-g. PubMed PMID: 8348610.

5. Zhou X, Sasaki H, Lowe L, Hogan BL, Kuehn MR. Nodal is a novel TGF-beta-like gene expressed in the mouse node during gastrulation. Nature. 1993;361(6412):543-7. Epub 1993/02/11. doi: 10.1038/361543a0. PubMed PMID: 8429908.

6. Conlon FL, Lyons KM, Takaesu N, Barth KS, Kispert A, Herrmann B, et al. A primary requirement for nodal in the formation and maintenance of the primitive streak in the mouse. Development. 1994;120(7):1919–28. Epub 1994/07/01. PubMed PMID: 7924997.

7. Feldman B, Gates MA, Egan ES, Dougan ST, Rennebeck G, Sirotkin HI, et al. Zebrafish organizer development and germ-layer formation require nodal-related signals. Nature. 1998;395(6698):181-5. Epub 1998/09/23. doi: 10.1038/26013. PubMed PMID: 9744277.

8. Birsoy B, Kofron M, Schaible K, Wylie C, Heasman J. Vg 1 is an essential signaling molecule in Xenopus development. Development. 2006;133(1):15–20. Epub 2005/11/26. doi: 10.1242/dev.02144. PubMed PMID: 16308332.

9. Bisgrove BW, Su YC, Yost HJ. Maternal Gdf3 is an obligatory cofactor in Nodal signaling for embryonic axis formation in zebrafish. Elife. 2017;6. Epub 2017/11/16. doi: 10.7554/eLife.28534. PubMed PMID: 29140249; PubMed Central PMCID: PMC5745076.

10. Montague TG, Schier AF. Vg1-Nodal heterodimers are the endogenous inducers of mesendoderm. Elife. 2017;6. Epub 2017/11/16. doi: 10.7554/eLife.28183. PubMed PMID: 29140251; PubMed Central PMCID: PMC5745085.

11. Pelliccia JL, Jindal GA, Burdine RD. Gdf3 is required for robust Nodal signaling during germ layer formation and left-right patterning. Elife. 2017;6. Epub 2017/11/16. doi: 10.7554/eLife.28635. PubMed PMID: 29140250; PubMed Central PMCID: PMC5745080.

12. Levin M, Johnson RL, Stern CD, Kuehn M, Tabin C. A molecular pathway determining left-right asymmetry in chick embryogenesis. Cell. 1995;82(5):803–14. Epub 1995/09/08. doi: 10.1016/0092-8674(95)90477-8. PubMed PMID: 7671308.

13. Collignon J, Varlet I, Robertson EJ. Relationship between asymmetric nodal expression and the direction of embryonic turning. Nature. 1996;381(6578):155-8. Epub 1996/05/09. doi: 10.1038/381155a0. PubMed PMID: 8610012.

14. Lowe LA, Supp DM, Sampath K, Yokoyama T, Wright CV, Potter SS, et al. Conserved left-right asymmetry of nodal expression and alterations in murine situs inversus. Nature. 1996;381(6578):158-61. Epub 1996/05/09. doi: 10.1038/381158a0. PubMed PMID: 8610013.

15. Pagan-Westphal SM, Tabin CJ. The transfer of left-right positional information during chick embryogenesis. Cell. 1998;93(1):25–35. Epub 1998/04/18. doi: 10.1016/s0092-8674(00)81143-5. PubMed PMID: 9546389.

16. Long S, Ahmad N, Rebagliati M. The zebrafish nodal-related gene southpaw is required for visceral and diencephalic left-right asymmetry. Development. 2003;130(11):2303–16. Epub 2003/04/19. doi: 10.1242/dev.00436. PubMed PMID: 12702646.

17. Montague TG, Gagnon JA, Schier AF. Conserved regulation of Nodal-mediated left-right patterning in zebrafish and mouse. Development. 2018;145(24). Epub 2018/11/18. doi: 10.1242/dev.171090. PubMed PMID: 30446628; PubMed Central PMCID: PMC6307886.

18. Gritsman K, Zhang J, Cheng S, Heckscher E, Talbot WS, Schier AF. The EGF-CFC protein one-eyed pinhead is essential for nodal signaling. Cell. 1999;97(1):121–32. Epub 1999/04/13. PubMed PMID: 10199408.

19. Yeo C, Whitman M. Nodal signals to Smads through Cripto-dependent and Cripto-independent mechanisms. Mol Cell. 2001;7(5):949–57. Epub 2001/06/08. doi: 10.1016/s1097-2765(01)00249-0. PubMed PMID: 11389842.

20. Cheng SK, Olale F, Bennett JT, Brivanlou AH, Schier AF. EGF-CFC proteins are essential coreceptors for the TGF-beta signals Vg1 and GDF1. Genes Dev. 2003;17(1):31–6. Epub 2003/01/07. doi: 10.1101/gad.1041203. PubMed PMID: 12514096; PubMed Central PMCID: PMC195969.

21. Baker JC, Harland RM. A novel mesoderm inducer, Madr2, functions in the activin signal transduction pathway. Genes Dev. 1996;10(15):1880–9. Epub 1996/08/01. doi: 10.1101/gad.10.15.1880. PubMed PMID: 8756346.

22. Hinck AP, Mueller TD, Springer TA. Structural Biology and Evolution of the TGF-beta Family. Cold Spring Harb Perspect Biol. 2016;8(12). Epub 2016/09/18. doi: 10.1101/cshperspect.a022103. PubMed PMID: 27638177; PubMed Central PMCID: PMC5131774.

23. Miyazono K, Olofsson A, Colosetti P, Heldin CH. A role of the latent TGF-beta 1-binding protein in the assembly and secretion of TGF-beta 1. EMBO J. 1991;10(5):1091–101. Epub 1991/05/01. PubMed PMID: 2022183; PubMed Central PMCID: PMC452762.

24. Beck S, Le Good JA, Guzman M, Ben Haim N, Roy K, Beermann F, et al. Extraembryonic proteases regulate Nodal signalling during gastrulation. Nat Cell Biol. 2002;4(12):981–5. Epub 2002/11/26. doi: 10.1038/ncb890. PubMed PMID: 12447384.

25. Blanchet MH, Le Good JA, Mesnard D, Oorschot V, Baflast S, Minchiotti G, et al. Cripto recruits Furin and PACE4 and controls Nodal trafficking during proteolytic maturation. EMBO J. 2008;27(19):2580–91. Epub 2008/09/06. doi: 10.1038/emboj.2008.174. PubMed PMID: 18772886; PubMed Central PMCID: PMC2567404.

26. Seidah NG, Prat A. The biology and therapeutic targeting of the proprotein convertases. Nat Rev Drug Discov. 2012;11(5):367–83. Epub 2012/06/12. doi: 10.1038/nrd3699. PubMed PMID: 22679642.

27. Dohrmann CE, Kessler DS, Melton DA. Induction of axial mesoderm by zDVR-1, the zebrafish orthologue of Xenopus Vg1. Dev Biol. 1996;175(1):108–17. Epub 1996/04/10. doi: 10.1006/dbio.1996.0099. PubMed PMID: 8608857.

28. Reddy PS, Corley RB. Assembly, sorting, and exit of oligomeric proteins from the endoplasmic reticulum. Bioessays. 1998;20(7):546–54. Epub 1998/09/02. doi: 10.1002/(SICI)1521-1878(199807)20:7<546::AID-BIES5>3.0.CO;2-I. PubMed PMID: 9723003.

29. Wilkinson B, Gilbert HF. Protein disulfide isomerase. Biochim Biophys Acta. 2004;1699(1-2):35–44. Epub 2004/05/26. doi: 10.1016/j.bbapap.2004.02.017. PubMed PMID: 15158710.

30. Helenius A, Aebi M. Roles of N-linked glycans in the endoplasmic reticulum. Annu Rev Biochem. 2004;73:1019–49. Epub 2004/06/11. doi: 10.1146/annurev.biochem.73.011303.073752. PubMed PMID: 15189166.

31. Hammond C, Braakman I, Helenius A. Role of N-linked oligosaccharide recognition, glucose trimming, and calnexin in glycoprotein folding and quality control. Proc Natl Acad Sci U S A. 1994;91(3):913–7. Epub 1994/02/01. doi: 10.1073/pnas.91.3.913. PubMed PMID: 8302866; PubMed Central PMCID: PMC521423.

32. Flynn GC, Chappell TG, Rothman JE. Peptide binding and release by proteins implicated as catalysts of protein assembly. Science. 1989;245(4916):385-90. Epub 1989/07/28. doi: 10.1126/science.2756425. PubMed PMID: 2756425.

33. Flynn GC, Pohl J, Flocco MT, Rothman JE. Peptide-binding specificity of the molecular chaperone BiP. Nature. 1991;353(6346):726-30. Epub 1991/10/24. doi: 10.1038/353726a0. PubMed PMID: 1834945.

34. Blond-Elguindi S, Cwirla SE, Dower WJ, Lipshutz RJ, Sprang SR, Sambrook JF, et al. Affinity panning of a library of peptides displayed on bacteriophages reveals the binding specificity of BiP. Cell. 1993;75(4):717–28. Epub 1993/11/19. doi: 10.1016/0092-8674(93)90492-9. PubMed PMID: 7902213.

35. Tannahill D, Melton DA. Localized synthesis of the Vg1 protein during early Xenopus development. Development. 1989;106(4):775–85. Epub 1989/08/01. PubMed PMID: 2562668.

36. Waugh DS. An overview of enzymatic reagents for the removal of affinity tags. Protein Expr Purif. 2011;80(2):283–93. Epub 2011/08/30. doi: 10.1016/j.pep.2011.08.005. PubMed PMID: 21871965; PubMed Central PMCID: PMC3195948.

37. Wylie SJ, Adams M, Chalam C, Kreuze J, Lopez-Moya JJ, Ohshima K, et al. ICTV Virus Taxonomy Profile: Potyviridae. J Gen Virol. 2017;98(3):352–4. Epub 2017/04/04. doi: 10.1099/jgv.0.000740. PubMed PMID: 28366187; PubMed Central PMCID: PMC5797945.

38. Feng S, Sekine S, Pessino V, Li H, Leonetti MD, Huang B. Improved split fluorescent proteins for endogenous protein labeling. Nat Commun. 2017;8(1):370. Epub 2017/08/31. doi: 10.1038/s41467-017-00494-8. PubMed PMID: 28851864; PubMed Central PMCID: PMC5575300.

39. Cabrita LD, Gilis D, Robertson AL, Dehouck Y, Rooman M, Bottomley SP. Enhancing the stability and solubility of TEV protease using in silico design. Protein Sci. 2007;16(11):2360–7. Epub 2007/10/02. doi: 10.1110/ps.072822507. PubMed PMID: 17905838; PubMed Central PMCID: PMC2211701.

40. Cesaratto F, Lopez-Requena A, Burrone OR, Petris G. Engineered tobacco etch virus (TEV) protease active in the secretory pathway of mammalian cells. J Biotechnol. 2015;212:159–66. Epub 2015/09/04. doi: 10.1016/j.jbiotec.2015.08.026. PubMed PMID: 26327323.

41. Dale L, Matthews G, Tabe L, Colman A. Developmental expression of the protein product of Vg1, a localized maternal mRNA in the frog Xenopus laevis. EMBO J. 1989;8(4):1057–65. Epub 1989/04/01. PubMed PMID: 2519512; PubMed Central PMCID: PMC400914.

42. Chen C, Ware SM, Sato A, Houston-Hawkins DE, Habas R, Matzuk MM, et al. The Vg1-related protein Gdf3 acts in a Nodal signaling pathway in the pre-gastrulation mouse embryo. Development. 2006;133(2):319–29. Epub 2005/12/22. doi: 10.1242/dev.02210. PubMed PMID: 16368929.

43. Munro S, Pelham HR. A C-terminal signal prevents secretion of luminal ER proteins. Cell. 1987;48(5):899–907. Epub 1987/03/13. doi: 10.1016/0092-8674(87)90086-9. PubMed PMID: 3545499.

44. Jumper J, Evans R, Pritzel A, Green T, Figurnov M, Ronneberger O, et al. Highly accurate protein structure prediction with AlphaFold. Nature. 2021;596(7873):583-9. Epub 20210715. doi: 10.1038/s41586-021-03819-2. PubMed PMID: 34265844; PubMed Central PMCID: PMC8371605.

45. Haas IG, Wabl M. Immunoglobulin heavy chain binding protein. Nature. 1983;306(5941):387-9. Epub 1983/11/24. doi: 10.1038/306387a0. PubMed PMID: 6417546.

46. Bole DG, Hendershot LM, Kearney JF. Posttranslational association of immunoglobulin heavy chain binding protein with nascent heavy chains in nonsecreting and secreting hybridomas. J Cell Biol. 1986;102(5):1558–66. Epub 1986/05/01. doi: 10.1083/jcb.102.5.1558. PubMed PMID: 3084497; PubMed Central PMCID: PMC2114236.

47. Tu L, Sun TT, Kreibich G. Specific heterodimer formation is a prerequisite for uroplakins to exit from the endoplasmic reticulum. Mol Biol Cell. 2002;13(12):4221–30. Epub 2002/12/12. doi: 10.1091/mbc.e02-04-0211. PubMed PMID: 12475947; PubMed Central PMCID: PMC138628.

48. Hu CC, Bachmann T, Zhou G, Liang FX, Ghiso J, Kreibich G, et al. Assembly of a membrane receptor complex: roles of the uroplakin II prosequence in regulating uroplakin bacterial receptor oligomerization. Biochem J. 2008;414(2):195–203. Epub 2008/05/17. doi: 10.1042/BJ20080550. PubMed PMID: 18481938; PubMed Central PMCID: PMC4048928.

49. Sitia R, Neuberger M, Alberini C, Bet P, Fra A, Valetti C, et al. Developmental regulation of IgM secretion: the role of the carboxy-terminal cysteine. Cell. 1990;60(5):781–90. Epub 1990/03/09. doi: 10.1016/0092-8674(90)90092-s. PubMed PMID: 2107027.

50. Alberini CM, Bet P, Milstein C, Sitia R. Secretion of immunoglobulin M assembly intermediates in the presence of reducing agents. Nature. 1990;347(6292):485-7. Epub 1990/10/04. doi: 10.1038/347485a0. PubMed PMID: 2120591.

51. Melnick J, Dul JL, Argon Y. Sequential interaction of the chaperones BiP and GRP94 with immunoglobulin chains in the endoplasmic reticulum. Nature. 1994;370(6488):373-5. doi: 10.1038/370373a0. PubMed PMID: 7913987.

52. Mayer M, Kies U, Kammermeier R, Buchner J. BiP and PDI cooperate in the oxidative folding of antibodies in vitro. J Biol Chem. 2000;275(38):29421–5. Epub 2000/07/14. doi: 10.1074/jbc.M002655200. PubMed PMID: 10893409.

53. Meunier L, Usherwood YK, Chung KT, Hendershot LM. A subset of chaperones and folding enzymes form multiprotein complexes in endoplasmic reticulum to bind nascent proteins. Mol Biol Cell. 2002;13(12):4456–69. Epub 2002/12/12. doi: 10.1091/mbc.e02-05-0311. PubMed PMID: 12475965; PubMed Central PMCID: PMC138646.

54. Shen Y, Hendershot LM. ERdj3, a stress-inducible endoplasmic reticulum DnaJ homologue, serves as a cofactor for BiP’s interactions with unfolded substrates. Mol Biol Cell. 2005;16(1):40–50. Epub 20041103. doi: 10.1091/mbc.e04-05-0434. PubMed PMID: 15525676; PubMed Central PMCID: PMC539150.

55. Feige MJ, Groscurth S, Marcinowski M, Shimizu Y, Kessler H, Hendershot LM, et al. An unfolded CH1 domain controls the assembly and secretion of IgG antibodies. Mol Cell. 2009;34(5):569–79. Epub 2009/06/16. doi: 10.1016/j.molcel.2009.04.028. PubMed PMID: 19524537; PubMed Central PMCID: PMC2908990.

56. Marcinowski M, Höller M, Feige MJ, Baerend D, Lamb DC, Buchner J. Substrate discrimination of the chaperone BiP by autonomous and cochaperone-regulated conformational transitions. Nat Struct Mol Biol. 2011;18(2):150–8. Epub 20110109. doi: 10.1038/nsmb.1970. PubMed PMID: 21217698.

57. Haas IG. BiP (GRP78), an essential hsp70 resident protein in the endoplasmic reticulum. Experientia. 1994;50(11-12):1012–20. doi: 10.1007/BF01923455. PubMed PMID: 7988659.

58. Hendershot L, Wei J, Gaut J, Melnick J, Aviel S, Argon Y. Inhibition of immunoglobulin folding and secretion by dominant negative BiP ATPase mutants. Proc Natl Acad Sci U S A. 1996;93(11):5269–74. doi: 10.1073/pnas.93.11.5269. PubMed PMID: 8643565; PubMed Central PMCID: PMC39234.

59. Vanhove M, Usherwood YK, Hendershot LM. Unassembled Ig heavy chains do not cycle from BiP in vivo but require light chains to trigger their release. Immunity. 2001;15(1):105–14. Epub 2001/08/04. doi: 10.1016/s1074-7613(01)00163-7. PubMed PMID: 11485742.

60. Hendershot L, Bole D, Köhler G, Kearney JF. Assembly and secretion of heavy chains that do not associate posttranslationally with immunoglobulin heavy chain-binding protein. J Cell Biol. 1987;104(3):761–7. doi: 10.1083/jcb.104.3.761. PubMed PMID: 3102505; PubMed Central PMCID: PMC2114523.

61. Shusta EV, Raines RT, Pluckthun A, Wittrup KD. Increasing the secretory capacity of Saccharomyces cerevisiae for production of single-chain antibody fragments. Nat Biotechnol. 1998;16(8):773–7. Epub 1998/08/14. doi: 10.1038/nbt0898-773. PubMed PMID: 9702778.

62. Blanchet MH, Le Good JA, Oorschot V, Baflast S, Minchiotti G, Klumperman J, et al. Cripto localizes Nodal at the limiting membrane of early endosomes. Sci Signal. 2008;1(45):ra13. Epub 2008/11/13. doi: 10.1126/scisignal.1165027. PubMed PMID: 19001664.

63. Tessadori F, Noel ES, Rens EG, Magliozzi R, Evers-van Gogh IJ, Guardavaccaro D, et al. Nodal signaling range is regulated by proprotein convertase-mediated maturation. Dev Cell. 2015;32(5):631–9. Epub 2015/02/17. doi: 10.1016/j.devcel.2014.12.014. PubMed PMID: 25684355.

64. Ben-Haim N, Lu C, Guzman-Ayala M, Pescatore L, Mesnard D, Bischofberger M, et al. The nodal precursor acting via activin receptors induces mesoderm by maintaining a source of its convertases and BMP4. Dev Cell. 2006;11(3):313–23. Epub 2006/09/05. doi: 10.1016/j.devcel.2006.07.005. PubMed PMID: 16950123.

65. Meeker ND, Hutchinson SA, Ho L, Trede NS. Method for isolation of PCR-ready genomic DNA from zebrafish tissues. Biotechniques. 2007;43(5):610, 2, 4. Epub 2007/12/13. doi: 10.2144/000112619. PubMed PMID: 18072590.

66. Westerfield M. The zebrafish book: a guide for the laboratory use of zebrafish (Danio rerio). 4th ed: University of Oregon Press, Eugene; 2000.

67. Kimmel CB, Ballard WW, Kimmel SR, Ullmann B, Schilling TF. Stages of embryonic development of the zebrafish. Dev Dyn. 1995;203(3):253–310. Epub 1995/07/01. doi: 10.1002/aja.1002030302. PubMed PMID: 8589427.

68. Gibson DG, Young L, Chuang RY, Venter JC, Hutchison CA, 3rd, Smith HO. Enzymatic assembly of DNA molecules up to several hundred kilobases. Nat Methods. 2009;6(5):343–5. Epub 2009/04/14. doi: 10.1038/nmeth.1318. PubMed PMID: 19363495.

69. Muller P, Rogers KW, Jordan BM, Lee JS, Robson D, Ramanathan S, et al. Differential diffusivity of Nodal and Lefty underlies a reaction-diffusion patterning system. Science. 2012;336(6082):721-4. Epub 2012/04/14. doi: 10.1126/science.1221920. PubMed PMID: 22499809; PubMed Central PMCID: PMC3525670.

70. Pedelacq JD, Cabantous S, Tran T, Terwilliger TC, Waldo GS. Engineering and characterization of a superfolder green fluorescent protein. Nat Biotechnol. 2006;24(1):79–88. Epub 2005/12/22. doi: 10.1038/nbt1172. PubMed PMID: 16369541.

71. Kapust RB, Tozser J, Fox JD, Anderson DE, Cherry S, Copeland TD, et al. Tobacco etch virus protease: mechanism of autolysis and rational design of stable mutants with wild-type catalytic proficiency. Protein Eng. 2001;14(12):993–1000. Epub 2002/01/26. doi: 10.1093/protein/14.12.993. PubMed PMID: 11809930.

72. Nallamsetty S, Kapust RB, Tozser J, Cherry S, Tropea JE, Copeland TD, et al. Efficient site-specific processing of fusion proteins by tobacco vein mottling virus protease in vivo and in vitro. Protein Expr Purif. 2004;38(1):108–15. Epub 2004/10/13. doi: 10.1016/j.pep.2004.08.016. PubMed PMID: 15477088.

73. Raran-Kurussi S, Waugh DS. A dual protease approach for expression and affinity purification of recombinant proteins. Anal Biochem. 2016;504:30–7. Epub 2016/04/24. doi: 10.1016/j.ab.2016.04.006. PubMed PMID: 27105777; PubMed Central PMCID: PMC4877217.

74. Skala W, Goettig P, Brandstetter H. Do-it-yourself histidine-tagged bovine enterokinase: a handy member of the protein engineer’s toolbox. J Biotechnol. 2013;168(4):421–5. Epub 2013/11/05. doi: 10.1016/j.jbiotec.2013.10.022. PubMed PMID: 24184090; PubMed Central PMCID: PMC3863954.

75. Thisse C, Thisse B. High-resolution in situ hybridization to whole-mount zebrafish embryos. Nat Protoc. 2008;3(1):59–69. doi: 10.1038/nprot.2007.514. PubMed PMID: 18193022.

76. Rogers KW, Lord ND, Gagnon JA, Pauli A, Zimmerman S, Aksel DC, et al. Nodal patterning without Lefty inhibitory feedback is functional but fragile. Elife. 2017;6. Epub 2017/12/08. doi: 10.7554/eLife.28785. PubMed PMID: 29215332; PubMed Central PMCID: PMC5720593.

77. Schindelin J, Arganda-Carreras I, Frise E, Kaynig V, Longair M, Pietzsch T, et al. Fiji: an open-source platform for biological-image analysis. Nat Methods. 2012;9(7):676–82. Epub 2012/06/30. doi: 10.1038/nmeth.2019. PubMed PMID: 22743772; PubMed Central PMCID: PMC3855844.

